# Molecular anatomy of PLK1 master docking motifs

**DOI:** 10.1101/2025.03.28.645803

**Authors:** Long Ren, Raphael Gasper, Marion E. Pesenti, Petra Janning, Franziska Müller, Carolin Koerner, Petra Geue, Sabine Wohlgemuth, Arianna Esposito Verza, Ingrid R. Vetter, Andrea Musacchio

**Affiliations:** Department of Mechanistic Cell Biology, Max Planck Institute of Molecular Physiology, Otto-Hahn-Straße 11, 44227 Dortmund, Germany; Crystallography and Biophysics Facility, Max Planck Institute of Molecular Physiology, Otto-Hahn-Straße 11, 44227 Dortmund, Germany; Mass Spectrometry Core Facility, Max Planck Institute of Molecular Physiology, Otto-Hahn-Straße 11, 44227 Dortmund, Germany; Centre for Medical Biotechnology, Faculty of Biology, University Duisburg-Essen, Essen, Germany

**Keywords:** PLK1, PBD, CENP-U, kinase, mitosis, CCAN, kinetochore, CDK1, phosphorylation, structure

## Abstract

The Polo-box domain (PBD) localizes Polo-like kinase 1 (PLK1) near mitotic substrates required for chromosome biorientation. Recent work on mitotic kinetochores showed PLK1 docking begins hierarchically at master docking motifs on BUB1 and CENP-U. Whether master docking motifs have common molecular features remains poorly understood. Presence on CENP-U of two neighbouring motifs generated by initial CDK1 priming and subsequent PLK1 phosphorylation led us to hypothesize PBD dimerization might be involved. Using biochemical, biophysical, and modelling approaches, we gathered strong evidence that CENP-U contains a single master docking motif. The motif is very high affinity and sufficient to form extensive interactions with the PBD, engaging multiple pockets on its surface without obvious added benefits from dimerization. Comparisons with motifs in BUB1, BUBR1, and PRC1 suggests apparent commonalities of master PLK1 docking motifs. We discuss the implications of our observations for the mechanism of PLK1 activation.

## Introduction

The cell cycle, the universal process of division of a mother cell into two daughters, is the foundation of cellular life. In eukaryotes, this process is executed through master regulators named cyclin-dependent kinases (CDKs), whose progressive activation orders all crucial cell cycle events, from the replication of chromosomes in S-phase to their segregation from the mother into its two daughters during M-phase (mitosis) (Morgan, 2007). CDK activation ultimately also controls the activity of additional cell cycle regulators, among which are additional mitotic kinases (Saurin, 2018). A crucial mitotic kinase, Polo-like kinase 1 (PLK1), has emerged for its essential functions in a number of cell division events (Pintard and Archambault, 2018; Zitouni et al., 2014). For instance, PLK1 has been implicated in the regulation of spindle assembly, centrosome function, nuclear envelope breakdown, sister chromatid cohesion, kinetochore-microtubule interactions, spindle assembly checkpoint signalling, centromere propagation, and cytokinesis, among others (Nigg, 2001; Pintard and Archambault, 2018). How PLK1 controls these processes with exquisite spatial and temporal accuracy remains poorly understood.

Human PLK1 is a 603-residue protein consisting of an amino-terminal Ser/Thr catalytic domain (KD) separated through a 67-residue interdomain linker (IDL) and the Polo Cap (PC) helix from a carboxy-terminal polo-box domain (PBD) (Figure 1A) (Golsteyn et al., 1995; Holtrich et al., 1994; Mundt et al., 1997). The PBD consists of two polo box (PB) subdomains, PB1 and PB2 (residues 411-489 and 511-592, respectively), each with a preceding loop, named L1 and L2 respectively (Figure 1A) (Cheng et al., 2003; Elia et al., 2003b). Each PB consists of a six-stranded anti-parallel β-sheet followed by an α-helix (the β6α fold). The two PBs in a PBD form a tightly packed functional unit stabilized by multiple interactions within a shared hydrophobic core (Park et al., 2010).

**Figure 1.**
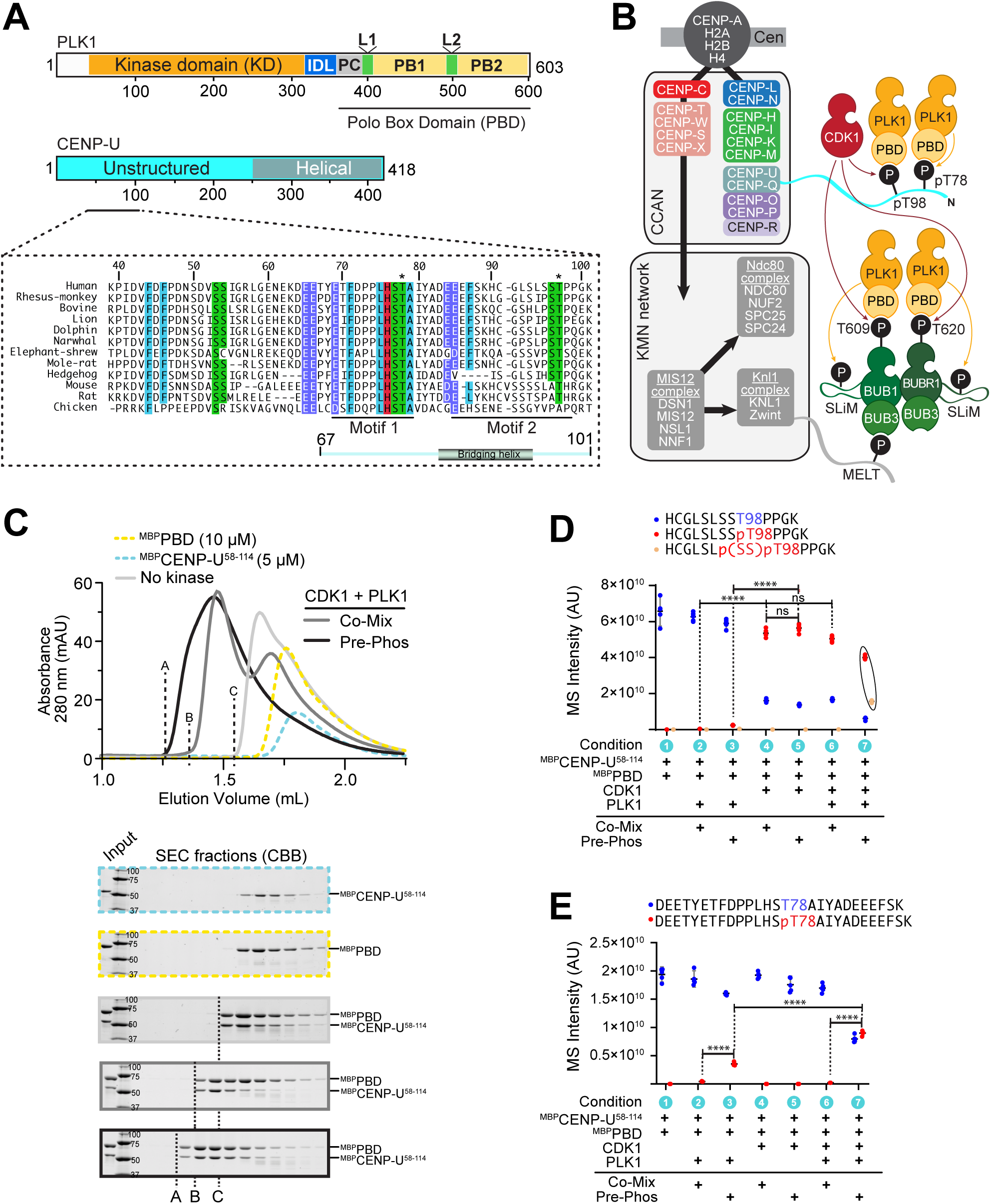
Assembly of a PLK1:CENP-U complex. (**A**) Schematic representation of the PLK1 and CENP-U. Abbreviations: KD = kinase domain; IDL = inter-domain linker; PC = Polo Cap; L1, L2 = Loop 1 and 2; PB1 and PB2 = Polo-box 1 and 2. (**B**) Hierarchical organization of the human kinetochore and localization domains of PLK1. CCAN = constitutive centromere associated network (inner kinetochore); KMH = KNL1 complex-MIS12 complex-NDC80 complex. Unstructured tail of CENP-U provides docking motifs for PLK1. Unstructured region of KNL1 provides phosphorylated motifs (MELTs) that recruit BUB3:BUB1, which in turn recruits BUB3:BUBR1. CDK1 phosphorylation of BUB1 and BUBR1 promotes recruitment of PLK1, which in turn contributes to phosphorylation of short linear motifs (SLiM) on BUB1 and BUBR1 that recruit PP2A phosphatase. (**C**) Analytical SEC profiles and corresponding SDS-PAGE for phosphorylation-dependent and independent binding of the PBD and CENP-U. A, B, and C indicate the elution front of different interacting species. Equivalent species will be labelled with the same letter throughout the paper. (**D**) Mass spectrometry analysis of phosphorylation of T98^CENP-U^ (Motif 2; CDK1) under co-mixing or pre-phosphorylation conditions. (**E**) Mass spectrometry analysis of T78^CENP-U^ (Motif 1; PLK1) under co-mixing or pre-phosphorylation. In panels D and E, data from five replicates were imported into GraphPad Prism 9.0 (GraphPad Software). Statistical analysis was performed using 2-way ANOVA test. Symbol indication: ns (not significant) = p > 0.05, ∗∗∗∗ = p ≤ 0.0001.

The interplay between the PLK1 KD and the PBD has attracted considerable interest and remains only partly understood. The PBD has at least two functions. First, it mediates the subcellular localization of PLK1 by binding to specific phosphorylated docking motifs in target proteins (Cheng et al., 2003; Elia et al., 2003a; Hanisch et al., 2006; Sharma et al., 2019; Singh et al., 2021). There is a quite strict requirement for targets to contain the sequence Ser-pThr-X or Ser-pSer-X (abbreviated respectively as S-pT-X or S-pS-X, where p indicates phosphorylation), with pThr preferred over pSer (Elia et al., 2003a). The X residue following pS/pT influences PBD binding selectivity only modestly, but influences greatly the choice of kinase introducing the phosphate. When X is Pro, the motif is a target of proline-directed kinases, like the CDKs. This widely documented mechanism, named “CDK priming” (an instantiation of nonself priming), links mitotic CDK activation to PLK1 localization and further PLK1 target phosphorylation (Elia et al., 2003a; Lowery et al., 2007). When X is not Pro, other kinases may be involved in the generation of PBD docking sites. “Self-priming” may occur when the phosphorylating kinase is PLK1 itself (Kang et al., 2011; Kang et al., 2006; Lee et al., 2008). Reflecting the preferences of the PLK1 kinase domain, this may occur especially in presence of Leu at position -3 relative to the target S/T, Asp, Asn, Glu, or Gln at position -2, and in absence of Pro at the +1 position (Alexander et al., 2011; Grosstessner-Hain et al., 2011; Santamaria et al., 2011).

Second, the PBD controls the activity of the KD through direct intramolecular interactions believed to dampen kinase activity (Jang et al., 2002; Lee and Erikson, 1997; Mundt et al., 1997; Xu et al., 2013). Extensive interactions between the PBD and the KD inhibit PLK1 kinase activity in the absence of phosphorylated ligands (Jang et al., 2002; Kachaner et al., 2014; Xu et al., 2013). The interface between the L1 loop and the kinase small lobe, and a sandwich of the interdomain-linker (IDL) between the kinase hinge domain and the L2 loop, are the main contacts (Xu et al., 2013). Peptide binding is believed to restructure and stabilize the L2 loop, arguably through the action of the phosphate group, disrupting these contacts and releasing the kinase active site from its partial occlusion in the auto-inhibited form (Chapagai et al., 2023; Park et al., 2023; Xu et al., 2013). Through these mechanisms, phosphorylated targets of the PBD may unleash kinase activity to promote phosphorylation of substrate locally, near the target site, eliciting sequential phosphorylation (Bertran et al., 2011; Lobjois et al., 2011; Matthess et al., 2014; O’Donovan et al., 2013; Zhang et al., 2009).

Importantly, the PBD appears to be dispensable for most mitotic functions of PLK1, but completion of chromosome biorientation requires it (Hanisch et al., 2006; Liu et al., 2011; Seong et al., 2002; Sharma et al., 2019). Work on the kinetochore, a large protein assembly on chromosomes that promotes microtubule binding and biorientation (Cheeseman and Desai, 2008; Musacchio and Desai, 2017), identified several kinetochore proteins as potential binding sites for the PLK1 PBD (Amin et al., 2014; Bel Borja et al., 2024; Geraghty et al., 2021; Goto et al., 2006; Kang et al., 2011; Kang et al., 2006; Kim et al., 2015; Kim et al., 2014; Lee et al., 2015; Maia et al., 2012; Matsumura et al., 2007; Nishino et al., 2006; Pouwels et al., 2014; Qi et al., 2006; Sun et al., 2012; Taylor et al., 2023; Yeh et al., 2013; Zhuo et al., 2015). Two of these, CENP-U (a.k.a. PBIP1) and BUB1, respectively residing in the inner and outer kinetochores (Figure 1B), emerged as master docking motifs for PLK1, as their suppression eliminated PLK1 recruitment to kinetochores altogether (Chen et al., 2021; Nguyen et al., 2021; Singh et al., 2021). Thus, recruitment of PLK1 to the kinetochore and other cellular locales may be hierarchical, with few high-affinity master binding motifs being essential for initial targeting and persistent activation of PLK1. This may be followed by local phosphorylation of other targets, in some cases also leading to the creation of secondary more transient docking sites. Recent work identifying a master docking motif in M18BP1 for recruitment of PLK1 to anaphase kinetochores is consistent with this concept (Conti et al., 2024; Parashara et al., 2024).

Understanding the mechanism of PLK1 activation is urgent also in view of evidence that this kinase, at least in its more fundamental state, appears to be rather poorly active, at least in comparison to other mitotic kinases (Johnson et al., 2008; Kothe et al., 2007). What defines master docking motifs, and whether self- or nonself-priming is preferentially involved in their generation, however, remains poorly understood. PLK1 recruitment to its putative master target motifs at the kinetochores, BUB1 and CENP-U, engaging both self- and nonself-priming mechanisms, exemplifies the potential molecular complexity of this problem. BUB1 binds PLK1 after CDK-mediated phosphorylation of T609 (Cordeiro et al., 2020; Corno et al., 2023; Houston et al., 2023; Ikeda and Tanaka, 2017; Jia et al., 2016; Qi et al., 2006; Singh et al., 2021). BUB1 also recruits BUBR1 (Overlack et al., 2015; Zhang et al., 2015), which itself interacts with PLK1 after CDK-mediated phosphorylation of T620. Recruitment of PLK1, in turn, facilitates binding of PP2A phosphatase to the kinetochore (Cordeiro et al., 2020; Corno et al., 2023; Elowe et al., 2007; Huang et al., 2008; Kruse et al., 2013; Matsumura et al., 2007; Song et al., 2024; Suijkerbuijk et al., 2012; Vallardi et al., 2017; Wong and Fang, 2007).

CENP-U, a 416-residue protein, contains a C*-* terminal helical region (residues 250-416) playing a structural role in the inner kinetochore, and an unstructured N*-*terminal segment (residues 1-249) of uncertain function (Figure 1A) (Singh et al., 2021; Yan et al., 2024). PLK1 phosphorylates T78 of CENP-U and subsequently binds to the phosphorylated motif, an example of self-priming (Kang et al., 2011; Kang et al., 2006; Lee et al., 2008). More recent evidence, however, suggested that T78 is inefficiently phosphorylated by PLK1 without prior phosphorylation of T98, a site within an S-T-P CDK1 substrate motif. The latter recruits PLK1 to enhance subsequent phosphorylation of T78, in a “relay priming” mechanism (Singh et al., 2021).

Initial biochemical experiments demonstrated that when T78 and T98 are concomitantly phosphorylated, two PBDs can dock side-a-side on CENP-U (Singh et al., 2021). Dimerization can be harnessed to increase the effective binding affinity to a target and is also a mechanism frequently used in allosteric kinase activation (Erlendsson and Teilum, 2020; Lavoie et al., 2014). Thus, the observations on CENP-U may reflect a strategy to promote local activation. Constellations of tandem PBD docking motifs seemingly similar to the one found in CENP-U are also present in BUB1, as well as in other PLK1 binding partners outside kinetochores (Singh et al., 2021). Nonetheless, the generality of this mechanism is unknown, and so are the exact determinants of high-affinity binding of the PBD to target sequences, a property plausibly required for their ability to qualify as master PLK1 docking motifs. MAP205, an established biochemical inhibitor of PLK1, engages the phosphoresidue-binding site using a phospho-mimetic strategy. It also extends further onto the PBD surface, interacting with a “cryptic pocket” previously shown to engage in recognition of a peptide encompassing residues 71-79 of CENP-U (Archambault et al., 2008; Kachaner et al., 2014; Liu et al., 2011; Sharma et al., 2019; Sledz et al., 2011; Xu et al., 2013).

Here, we revisited the mechanism of CDK priming on CENP-U and the generality of the PBD dimerization mechanism that emerged from initial analysis of the interaction of PLK1 with CENP-U (Singh et al., 2021). A crystal structure of two PBD domains bound to a CENP-U fragment encompassing both the pT78 and pT98 motifs revealed similarities and differences in the organization of the two bound PBDs. We confirm that initial CDK priming on T98 is required for efficient T78 relay-priming phosphorylation by PLK1. We show that PLK1 has extremely high affinity for the motif encompassing pT78, approximately 100-fold stronger than for the pT98 motif. The high binding affinity for the pT78 motif reflects an extended interaction interface that occupies various docking stations on the PB1 and the PB2. On the basis of these observation, quantitative analyses, and additional structural modelling, we propose that PLK1 master docking motifs, whether generated by CDK1 or PLK1 itself, occupy a much greater area of the PBD surface than hitherto realised, and have several common structural features that distinguish them from transient sites. Collectively, our results have important implications for understanding PLK1 function.

## Results

### pT98 primes phosphorylation of T78

We reconstituted *in vitro* a PLK1:CENP-U complex with PBD fused N-terminally to maltose-binding-protein (^MBP^PBD) and residues 58-114 of CENP-U fused N-terminally to MBP (^MBP^CENP-U^58-114^). In analytical size-exclusion chromatography (SEC), both ^MBP^CENP-U^58-114^ and ^MBP^PBD eluted in single peaks (Figure 1C, yellow and blue lines). When mixed at a molar ratio of 2:1 (10 µM ^MBP^PBD and 5 µM ^MBP^CENP-U^58-114^), the two species underwent a leftward shift in elution volume and co-eluted (Figure 1C; light grey line), suggesting that at the concentrations of these experiments complex formation can occur also in the absence of phosphorylation. Using non-overlapping constructs encompassing the first (T78) or second (T98) phosphorylation site and neighbouring residues (^MBP^CENP-U^58-83^ and ^MBP^CENP-U^84-114^, which we also refer to as Motif 1 and Motif 2, respectively), we determined that the PBD has residual binding affinity for unphosphorylated Motif 1, but not for unphosphorylated Motif 2 (Figure S1A-C and see below).

To phosphorylate T78 and T98 of ^MBP^CENP-U^58-114^, we added catalytic amounts (typically 1:30 molar ratio) of active CDK1 and PLK1 kinase (see Methods). Successful phosphorylation was confirmed by Pro-Q™ Diamond staining, which specifically stains phosphorylated protein. We tested either 1) pre-phosphorylating with both kinases ^MBP^CENP-U^58-114^ and then adding ^MBP^PBD for complex formation (pre-phosphorylation method); or 2) mixing ^MBP^CENP-U^58-114^ and ^MBP^PBD with both kinases for simultaneous phosphorylation and complex formation (co-mixing method). SEC analysis demonstrated that pre-phosphorylation (Figure 1C; black line) resulted in an apparently stoichiometric complex with smaller elution volume relative to the complex generated by co-mixing (dark grey line).

We used mass spectrometry (MS) to further assess CENP-U^58-114^ phosphorylation potential in the co-mixing and pre-phosphorylation regimes. CDK1 (used at 100 nM against 5 μM substrate) was sufficient for robust phosphorylation of T98, regardless of whether pre-phosphorylation or co-mixing were used (Figure 1D, conditions 4-5). Conversely, phosphorylation in presence of PLK1 (also used at 100 nM; Figure 1D, conditions 2-3) was either absent or below detection limit. Under the specific conditions used, PLK1 needed pre-phosphorylation to phosphorylate CENP-U on T78, with modest phosphorylation in isolation and efficient phosphorylation when combined to CDK1 (Figure 1E, condition 3 and 7). We surmise that significant binding affinity of the PBD for Motif 1 even before phosphorylation suppresses phosphorylation of T78 when reactions are carried out in presence of PBD in the co-mixing procedure. We do not attribute physiological significance to this observation, but discuss it as a relevant factor in the outcome of biochemical procedures to obtain the fully phosphorylated complex. As T98 is strongly phosphorylated even in absence of PLK1 (Figure 1D, conditions 4-5), these results imply that efficient phosphorylation of T78^CENP-U^ under the stringent conditions of this experiment only occurs when T98 is phosphorylated, i.e. when PLK1 and CDK1 are concomitantly present, confirming that CDK1 primes phosphorylation of T78 by PLK1 in a mechanism of “relay priming”. We note that despite a leucine at the -3 position, T78 has a less favourable histidine at the -2 position (see Introduction), rendering T78 a non-ideal PLK1 substrate and likely explaining why relay priming facilitates its phosphorylation.

### Crystallization and structure determination

We generated untagged versions of CENP-U^58-114^ and the PBD for crystallization using pre-phosphorylation (see Methods). Crystals were obtained as described in Methods and had properties summarized in Table 1. Structure determination to a minimal Bragg spacing of 2.15 Å was obtained using the molecular replacement method for phasing (Table 1, dataset 1). The CENP-U^58-114^ peptide bridges two sequentially positioned PBDs, each of which bound to one of the two phospho-Thr, for which there is unequivocal electron density (pT78 and pT98; Figure 2A and Figure S2A-B). A short helical segment of CENP-U, the bridging helix (residues E84 to L93), connects the PBD-binding motifs, buried at the interface between the two PBDs. The latter don’t make any direct contact in the complex.

**Table I.**
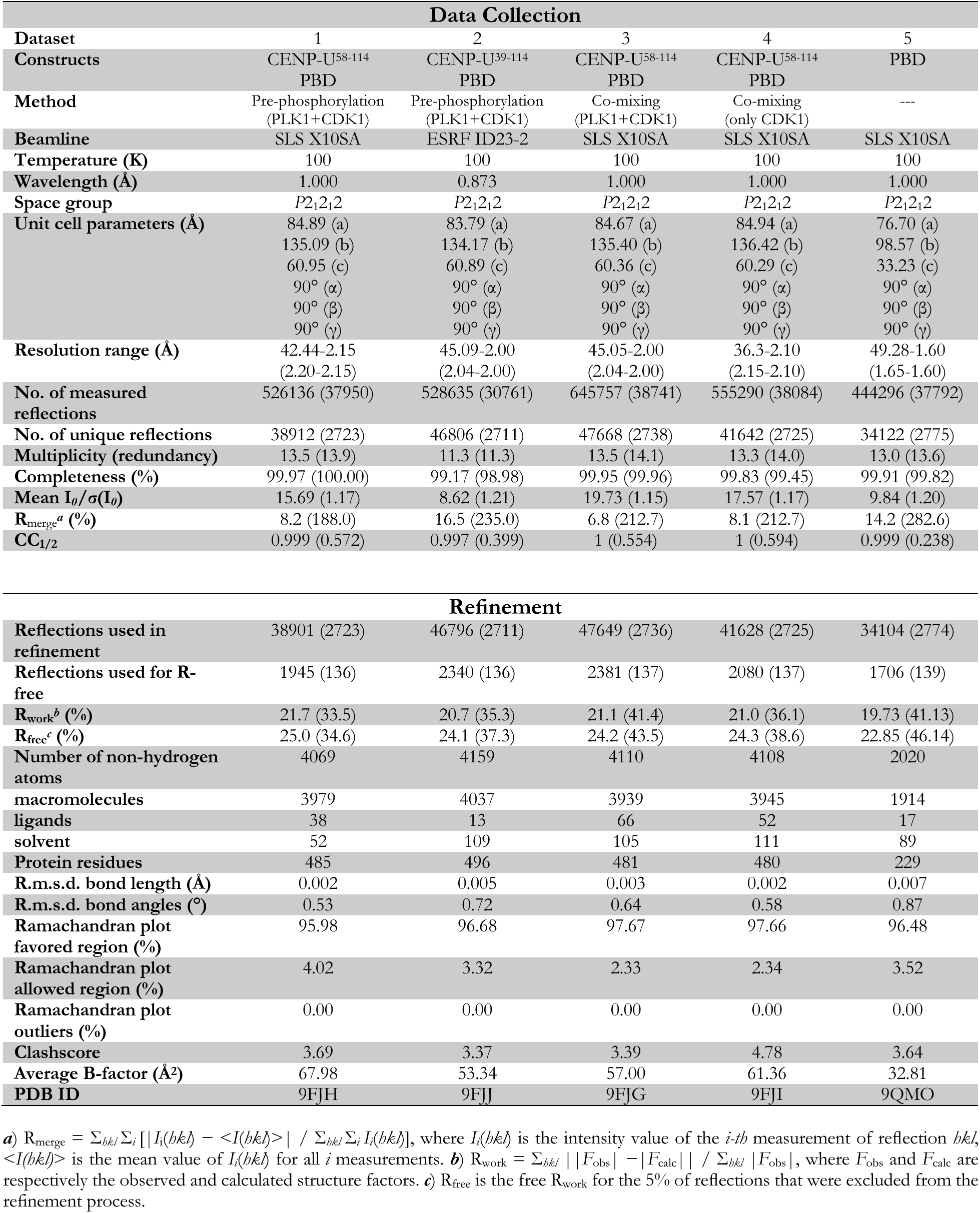
Data collection and refinement statistics.

**Figure 2.**
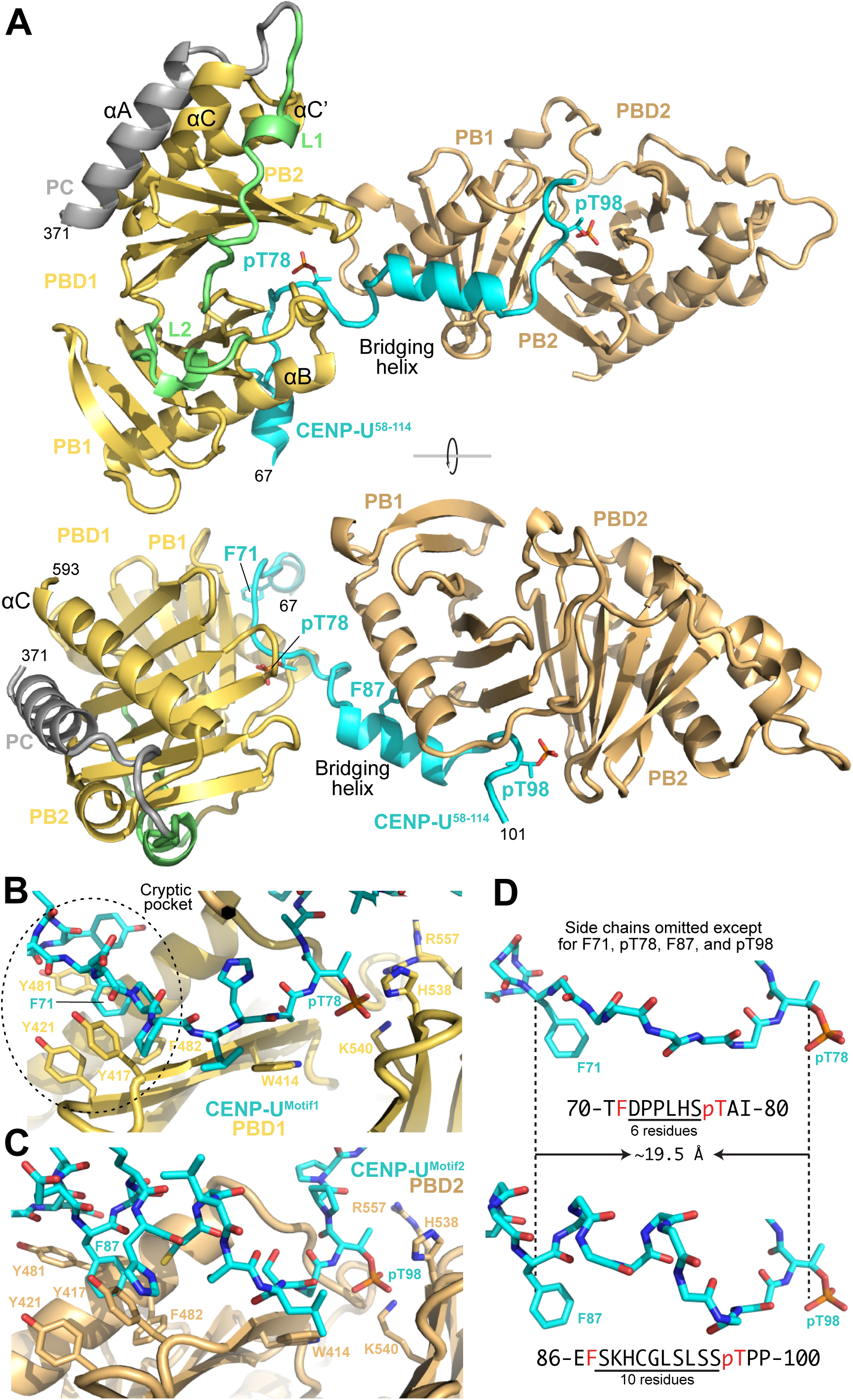
Structural basis for CENP-U binding by PLK1. (**A**) Two rotated cartoon models of the crystal structure from Dataset 1 of a trimeric complex formed by CENP-U^58-114^ with both T78^CENP-U^ and T98^CENP-U^ phosphorylated (cyan) and two bound PBDs (identified as PBD1 and PBD2 and shown in yelloworange and wheat, respectively). Key residues are indicated in stick representation. Colouring as per Figure 1A. Helices were named as in (Elia et al., 2003b). (**B**) Closeup of the pT78 motif represented in sticks against a cartoon of PBD1 with key residues highlighted in stick representation. The position of the cryptic pocket hosting F71 (and F87 in panel C) is highlighted. (**C**) As in panel B for pT98. (**D**) Comparison of Motif 1 and Motif 2 with Phe (F) and pT representing two pins of a staple inserted respectively into the hydrophobic ‘cryptic pocket’ and the electrostatic ‘pincer’ of PBD1. The distance between Cα is essentially identical.

Motifs 1 and 2 bind their cognate PBDs in apparently similar ways (Figure 2B-C). The side chains of the two phosphorylated residues, pT78^CENP-U^ and pT98^CENP-U^, are bound essentially identically through a “pincer” made by H538^PLK1^ and K540^PLK1^ (Cheng et al., 2003; Elia et al., 2003b), located in the second polo-box (PB2) of the PBD (Figure 2B-C). R557^PLK1^ (more evident at the binding site for pT78^CENP-U^) and a few water molecules complete the shell of interactions around the phosphate. In addition, both CENP-U motifs make extensive contacts with the PB1, most notably by inserting phenylalanine side chains (F71^CENP-U^ and F87^CENP-U^) into a cavity formed by various aromatic side chains, including those of Y417^PLK1^, Y421^PLK1^, Y481^PLK1^, and F482^PLK1^ (Figure 2B-C) and previously identified as the “cryptic pocket” (also identified as Tyr pocket) (Sharma et al., 2019; Sledz et al., 2011) for reasons that are clarified below. After mutating F87 to alanine, the stable complex of apparent 1:1 stoichiometry between ^MBP^CENP-U^84-114^ (encompassing Motif 2) and ^MBP^PBD obtained after CDK1 phosphorylation was shifted to a greater elution volume, indicative of a loss of stability and confirming the importance of this interaction (Figure S1D-E).

Despite different numbers of residues (six and ten, respectively), the distances between C_α_ atoms in the two F71^CENP-U^-pT78^CENP-U^ and F87^CENP-U^-pT98^CENP-U^ pairs are essentially identical (∼19.5 Å) (Figure 2D). Two consecutive prolines (P73-P74) promote a more extended main chain conformation between F71^CENP-U^ and pT78^CENP-U^, whereas the connection between F87^CENP-U^ and pT98^CENP-U^ is less extended and partly helical, explaining this. Thus, a common element of the two PLK1-binding motifs of CENP-U is that they bind the PBD as a staple, with an electrostatic interaction network around the phospho-threonine, and a hydrophobic interaction around a phenylalanine preceding the phospho-threonine, with an adaptable number of residues in between. Additional element that instead diversify these motifs are discussed below.

### Quantitative analysis of the CENP-U:PBD interaction

Our observations so far are that phosphorylation of T78 by PLK1 is facilitated by prior phosphorylation of T98 by CDK1, and that two PBD domains can be bound to these residues at the same time, confirming and extending our previous results (Singh et al., 2021). We previously hypothesized that dimerization of PLK1 on a target may give rise to a cooperative, more stable assembly, and that this could explain the ability of CENP-U to act as a master PLK1 docking site (Singh et al., 2021). Importantly, our new structure of two PBDs bound to CENP-U did not reveal obvious evidence of cooperativity, as the two PBD do not touch each other and are separated by the bridging helix. Therefore, we wanted to gain a more quantitative perspective of the mechanism of PLK1 binding to CENP-U.

For this, we measured binding thermodynamics and kinetics using biolayer interferometry (BLI) and isothermal titration calorimetry (ITC), obtaining association and dissociation rate constants (k_on_ and k^off^) and dissociation constant (K_D_). We began by characterizing the interaction with Motif 1 by BLI. To obtain T78 phosphorylation in absence of T98, we increased the concentration of CENP-U^58-114^ and active PLK1 kinase to bypass priming and phosphorylate T78 with only PLK1 (see Methods). Biotin-conjugated ^MBP^PBD immobilized on the instrument’s sensor tip surface bound to unphosphorylated ^MBP^CENP-U^58-83^ (only encompassing Motif 1) with a dissociation constant (K_D_) of 10 µM (Table 2a, entry 1; Figure S3A-L shows sensorgrams for all listed experiments). This relatively high affinity, and the fact that it arises from a very slow k_off_, explains the ability of the PBD to inhibit T78 phosphorylation in the co-mixing regime. PLK1-phosphorylated ^MBP^CENP-U^58-83^, on the other hand, bound with a K_D_ of 2.5 nM (Table 2a, entry 2). Thus, Motif 1 binds the PBD with very high binding affinity even in the absence of dimerization. Mutations of residues in Motif 1, including Y68A and F71A, that bind at or near the cryptic pocket and complement the interaction of pT78 at the pincer severely affected binding of ^MBP^PBD to Motif 1 (Table 2a, entry 3-4). An ^MBP^CENP-U^58-114^ construct carrying mutations impairing Motif 2, and phosphorylated with only PLK1, had a dissociation constant very similar to that observed with the phosphorylated Motif 1 (Table 2a, entry 5). Differently from Motif 1, unphosphorylated Motif 2 did not bind the PBD in BLI experiments, while a peptide phosphorylated on T98 bound with a K_D_ of ∼400 nM (Table 2a, entries 6-7).

**Table 2.** BLI analysis quantified binding kinetic constants.

| Table 2a \| Binding parameters of PBD to variants of CENP-U (unmodified or phosphorylated) |  |  |  |  |  |
| --- | --- | --- | --- | --- | --- |
| Entry | Immobilized Ligand | Analyte | K<sub>D</sub> (M) | k<sub>on</sub> (1/Ms) | k<sub>off</sub> (1/s) |
| 1 | MBPPBD | MBP<sub>CENP-U</sub><sup>58-83</sup> | 1.0 × 10<sup>-5</sup> | 2.8 × 10<sup>1</sup> | 2.8 × 10<sup>-4</sup> |
| 2 |  | MBP<sub>CENP-U</sub><sup>58-83/pT78</sup> | 2.5 × 10<sup>-9</sup> | 1.4 × 10<sup>5</sup> | 3.4 × 10<sup>-4</sup> |
| 3 |  | MBP<sub>CENP-U</sub><sup>58-83/F71A/pT78</sup> | 8.8 × 10<sup>-8</sup> | 8.7 × 10<sup>4</sup> | 7.7 × 10<sup>-3</sup> |
| 4 |  | MBP<sub>CENP-U</sub><sup>58-83/Y68A/F71A/pT78</sup> | 2.3 × 10<sup>-7</sup> | 4.7 × 10<sup>4</sup> | 1.1 × 10<sup>-2</sup> |
| 5 |  | MBP<sub>CENP-U</sub><sup>58-114/pT78/F87A/T98V</sup> | 7.2 × 10<sup>-9</sup> | 1.4 × 10<sup>5</sup> | 1.0 × 10<sup>-3</sup> |
| 6 |  | MBP<sub>CENP-U</sub><sup>84-114</sup> | N.B.<sup>a</sup> | --- | --- |
| 7 |  | MBP<sub>CENP-U</sub><sup>84-114/pT98</sup> | 4.0 × 10<sup>-7</sup> | 7.1 × 10<sup>3</sup> | 2.8 × 10<sup>-3</sup> |
| Table 2b \| Binding parameters of PBD or PLK1<sup>FL</sup> to variants of CENP-U (unmodified or phosphorylated) |  |  |  |  |  |
| 8 | MBP<sub>CENP-U</sub><sup>58-114/pT78</sup> | MBP<sub>PBD</sub> | 1.9 × 10<sup>-9</sup> | 2.8 × 10<sup>5</sup> | 5.2 × 10<sup>-4</sup> |
| 9 |  | 6His<sub>PLK1</sub><sup>FL</sup> | 2.2 × 10<sup>-8</sup> | 1.5 × 10<sup>5</sup> | 3.3 × 10<sup>-3</sup> |
| 10 | MBP<sub>CENP-U</sub><sup>58-114/pT98</sup> | MBP<sub>PBD</sub> | 1.6 × 10<sup>-8</sup> (K<sub>D1</sub>)<br>4.0 × 10<sup>-6</sup> (K<sub>D2</sub>) | 4.5 × 10<sup>4</sup><br>7.4 × 10<sup>3</sup> | 7.1 × 10<sup>-4</sup><br>2.9 × 10<sup>-2</sup> |
| 11 |  | 6His<sub>PLK1</sub><sup>FL</sup> | 4.2 × 10<sup>-6</sup> (K<sub>D1</sub>)<br>1.1 × 10<sup>-5</sup> (K<sub>D2</sub>) | 2.3 × 10<sup>3</sup><br>5.6 × 10<sup>4</sup> | 1.0 × 10<sup>-2</sup><br>6.0 × 10<sup>-1</sup> |
| 12 | MBP<sub>CENP-U</sub><sup>58-114/pT78/pT98</sup> | MBP<sub>PBD</sub> | 2.7 × 10<sup>-9</sup> (K<sub>D1</sub>)<br>1.0 × 10<sup>-6</sup> (K<sub>D2</sub>) | 4.5 × 10<sup>4</sup><br>1.8 × 10<sup>4</sup> | 1.2 × 10<sup>-4</sup><br>1.8 × 10<sup>-2</sup> |
| 13 |  | 6His<sub>PLK1</sub><sup>FL</sup> | 1.0 × 10<sup>-8</sup> (K<sub>D1</sub>)<br>1.6 × 10<sup>-6</sup> (K<sub>D2</sub>) | 1.7 × 10<sup>5</sup><br>9.4 × 10<sup>3</sup> | 1.8 × 10<sup>-3</sup><br>1.5 × 10<sup>-2</sup> |

<sup>a</sup>) N.B., no binding.

To investigate the putative cooperativity of the interaction of PBDs with CENP-U^58-114^, we biotin-conjugated and immobilized the latter and measured binding affinity to singly and doubly phosphorylated CENP-U. We reasoned that cooperative binding should manifest itself as a noticeable increase of binding affinity under conditions allowing the occupation of both binding sites by the PBD. We initially immobilized ^MBP^CENP-U^58-114/pT78^ and compared the binding affinity of the PBD with that of full length PLK1 (PLK1^FL^; Table 2b, entries 8-9). Fitting of binding kinetics indicated that ^MBP^CENP-U^58-114/pT78^ contains a single binding site (centred around Motif 1). This is consistent with the observation that the unphosphorylated Motif 2 does not bind the PBD (Table 2a, entry 6).

Next, we repeated the binding experiments on immobilized CENP-U^58-114/pT98^ (Table 2b, entries 10-11). In these two measurements, analysis of binding parameters suggested the existence of a primary, higher affinity binding site (K_D_1) and of a secondary, lower affinity binding site (K_D_2). In view of results in Table 2a, which were obtained with the immobilized PBD, K_D_1 likely reflects binding to the phosphorylated Motif 2, while K_D_2 reflects binding to the unphosphorylated Motif 1. Finally, we immobilized doubly phosphorylated ^MBP^CENP-U^58-114/pT78/pT98^ and once more, monitored binding of the PBD (Table 2b, entries 12-13). Also in this case, analysis of binding parameters suggested the existence of a primary, higher affinity binding site (K_D_1, presumably reflecting binding to Motif 1) and of a secondary, lower affinity binding site (K_D_2, presumably reflecting binding to Motif 2).

Using ITC, we measured the binding affinity of the PBD for singly phosphorylated pT78 and pT98 peptides. The ITC measurements were in excellent agreement to the BLI measurements, identifying a K_D_ of ∼380 nM for the pT98 peptide, and of ∼1.9 nM for the pT78 peptide (Figure S3M-O). These measurements, obtained with an approach orthogonal to BLI (ITC is performed in solution and measures the concentration of binding product at equilibrium, rather than assembly and disassembly kinetics), confirm that pT78 provides a very high affinity binding site for the PLK1 PBD. Collectively, several patterns emerge with clarity from these analyses. First, the PBD binds to pT78 with approximately 100-fold higher affinity than to pT98. Second, the PBD binds CENP-U with systematically higher affinity than the PLK1^FL^, quantifiable in approximately one order of magnitude difference in the dissociation constant. We confirmed this conclusion by using orthogonal approaches, including SEC (Figure S4A-D) and SEC-Multi Angle Light Scattering (SEC-MALS; Figure S5A-G). We can only speculate on the causes of this difference in binding affinity, but suspect it may reflect a) the larger hydrodynamic radius of PLK1^FL^ relative to the PBD, which in turn reflects on a larger k_off_; and b) an energy penalty associated with the disruption of intramolecular communication between the PBD and the kinase domain. Third, the presence of two PBDs on CENP-U does not appear to have a strong positive cooperative effect. If anything, presence of two phosphorylation sites may even appear to slightly increase the dissociation constants, i.e. act anti-cooperatively. From this, we conclude that dimerization, while clearly happening, may not be a crucial factor in the creation of a master site for PLK1 on CENP-U. Rather, our data suggest that CDK1 phosphorylates CENP-U on Thr98 to attract PLK1 to an otherwise poorly phosphorylated site, Thr78, to which PLK1 then binds extremely tightly, in line with the idea that pThr78 may represent a master PLK1 docking site.

### Determinants of master binding sites

As discussed in the Introduction, the precise molecular determinants of master docking sites for PLK1 recruitment remain unclear. To date, structural work has captured PBD interactions of short phosphopeptides binding exclusively in the vicinity of the phosphoresidue binding pocket (see for instance (Elia et al., 2003b; Garcia-Alvarez et al., 2007; Jia et al., 2015; Sledz et al., 2011; Yun et al., 2009)). In addition, structures of residues 71-79 of CENP-U in complex with the PBD identified an additional point of contact at the cryptic pocket (Sharma et al., 2019; Sledz et al., 2011). The structure calculated from Dataset 1 and our subsequent biochemical experiments add significantly to the picture. First, it appears that occupation of the cryptic pocket is more common than hitherto appreciated, as we see it not only with the pT78-containing motif, but also with the pT98-containing motif. Our BLI experiments showed reduced binding affinity when F71, which occupies the cryptic pocket, was mutated (Table 2). This is in line with previous experiments that reported an approximately 5-fold increase in binding affinity when this pocket becomes engaged (0.25 µM instead of 1.3 µM) (Sledz et al., 2011). Thus, occupation of the cryptic pocket leads to an increase of binding affinity relative to peptides with shorter N-terminal extensions.

Second, if the pT78-containing motif binds the PBD with at least 100-fold higher binding affinity even relative to the pT98-containing motif (Table 2) and to other previously tested binders, where measured affinities were generally in the high nanomolar to low micromolar range (Elia et al., 2003b; Garcia-Alvarez et al., 2007; Jia et al., 2015; Sledz et al., 2011; Song et al., 2024; Yun et al., 2009), additional interactions between the pT78 peptide and the PBD must be at stake, even beyond the cryptic pocket. A comparison of the two PBD:CENP-U sub-structures (Figure 3A) provides possible hints towards the source of this additional binding affinity. The CENP-U chain arriving from PBD1 contacts PBD2 immediately before the cryptic pocket (at Phe87). Conversely, there is extensive unexplored surface of PBD1 to which the CENP-U region preceding Phe71 could bind. Indeed, Tyr68 is an important determinant to the overall binding affinity of the CENP-U:PBD complex (Table 2), thus indicating that even residues preceding Phe71 contribute to increase the binding affinity of this motif for the PBD. Thus, by focusing on peptides targeting the pincer pocket, and occasionally the cryptic pocket, previous structural work on PBD:target interactions might have not captured the full interaction potential of the PBD.

**Figure 3.**
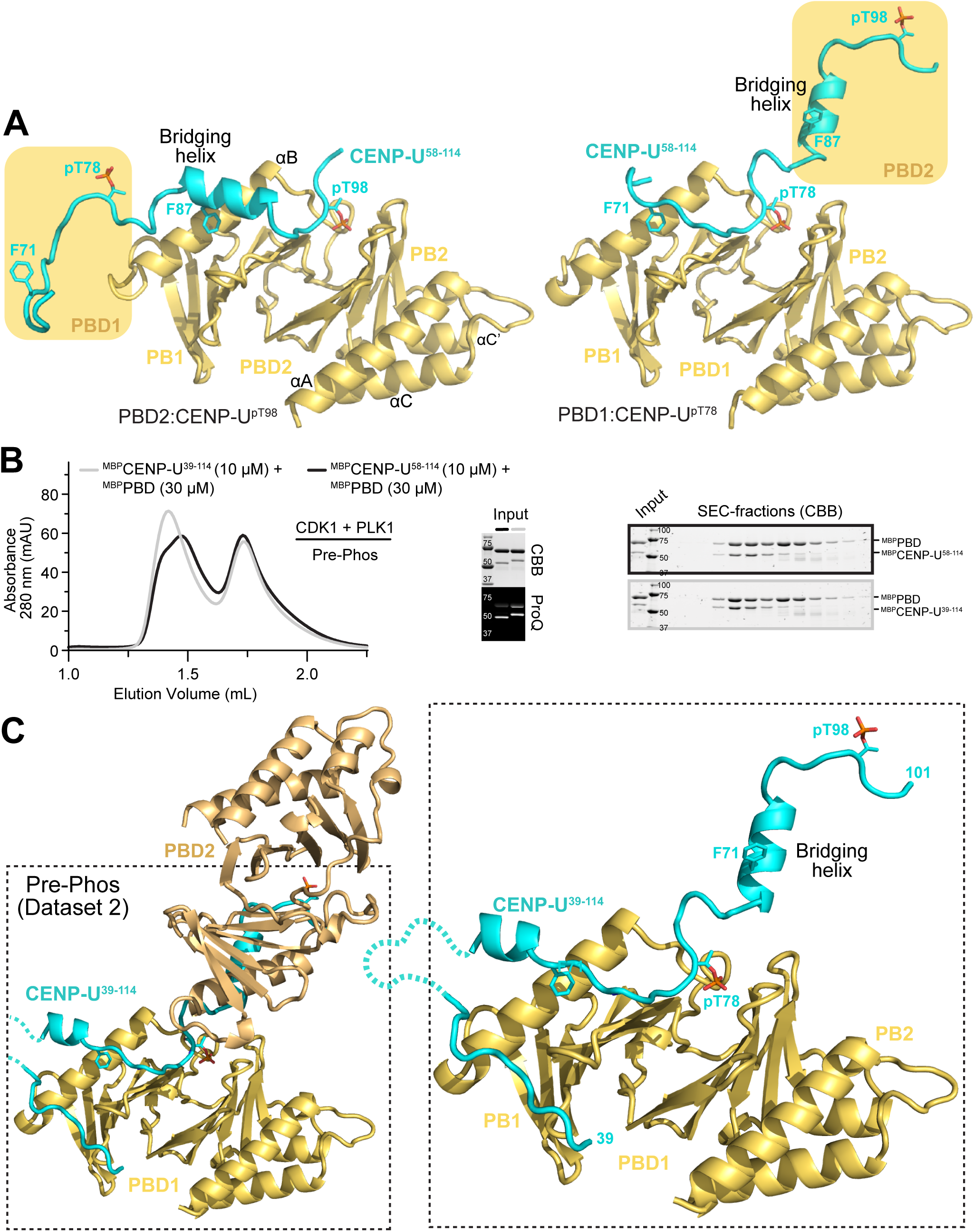
More extended interaction of CENP-U^39-114^. (**A**) Comparison of the interaction of CENP-U^58-114^ with PBD1 and PBD2 (right and left, respectively) suggests the region N-terminal to Motif 1 might interact with the exposed face of the PBD, like in the structure with MAP205 (see Figure 4F). (**B**) Comparison by SEC and SDS-PAGE of PBD complexes with CENP-U^58-114^ or CENP-U^39-114^ generated by pre-phosphorylation with CDK1 and PLK1. The ProQ^TM^ reagent interacts with phosphorylated species (see Methods). CBB = Coomassie Brilliant Blue. (**C**) Cartoon model of the crystal structure calculated from dataset 2 shows a very similar arrangement to that shown in Figure 2A (and identical colouring scheme), but with an N-terminal extension running transversally between the two PBs.

To further investigate this question, we extended the CENP-U construct on its N-terminal end by generating CENP-U^39-114^ and reconstituted its interaction with the PBD. In SEC experiments, the peak of the PBD:CENP-U^39-114^ complex eluted similarly to that of the PBD:CENP-U^58-114^ complex, but appeared more homogeneous (Figure 3B and Figure S6A). The PBD:CENP-U^39-114^ complex generated by pre-phosphorylation crystallized under conditions similar to those of the PBD:CENP-U^58-114^ complex, and we determined its 2.0 Å resolution crystal structure (Table 1, dataset 2; Figure 3C and Figure S6B-C). As expected, the structure of PBD:CENP-U^39-114^ is very similar to that of the PBD:CENP-U^58-114^ complex, with two PBDs bound consecutively to the CENP-U chain (Figure 3C). The main difference is that there is clear density for several residues in the N-terminal extension of CENP-U that were not included in the PBD:CENP-U^58-114^ complex, including a segment from residues 39 to 47 that appears to be stabilized in part by a crystal interface (Figure 3C and Figure S6B-C). Other parts of the segment were disordered and not visible in the electron density. Although not definitively proving that larger segments of the CENP-U N-terminal region contribute to the high binding affinity of motif 1 for the PBD, these observations strongly indicate the presence of additional contacts

### Predictions of extended interactions with the PBD

To complement the structural work, we performed an AlphaFold 3 (AF) (Abramson et al., 2024) prediction of a complex of the CENP-U N-terminal region with the PBD. AF predicted with high-confidence that the CENP-U N-terminal region meanders over the entire face of the PBD, anchoring first at a new exposed pocket in PB2, and then crossing over to PB1, making a prominent U-turn to interact with the cryptic pocket and with the pincer pocket (Figure 4A-C and Figure S7A). Equivalent high-confidence predictions for BUB1 and BUBR1 identified very similar binding modes (Figure 4D and Figure S7B-C). In all three cases, the first meaningful N-terminal contact of the target peptide is predicted to converge on the new pocket, where target peptides position Phe or Leu residues (Figure 4C). The pocket identified by AF in PB2 is formed by two highly conserved aromatics, F534 and Y582. Being positioned roughly pseudo-symmetrically to the cryptic pocket in PB1, we refer to it as the pseudo-cryptic pocket (Figure 4E). In addition, S439 and R441, positioned in PB1 but projecting their side chains towards PB2, are equivalent to the pincer residues H538 and K540 in PB2, and are also highly conserved (Figure 4E).

**Figure 4.**
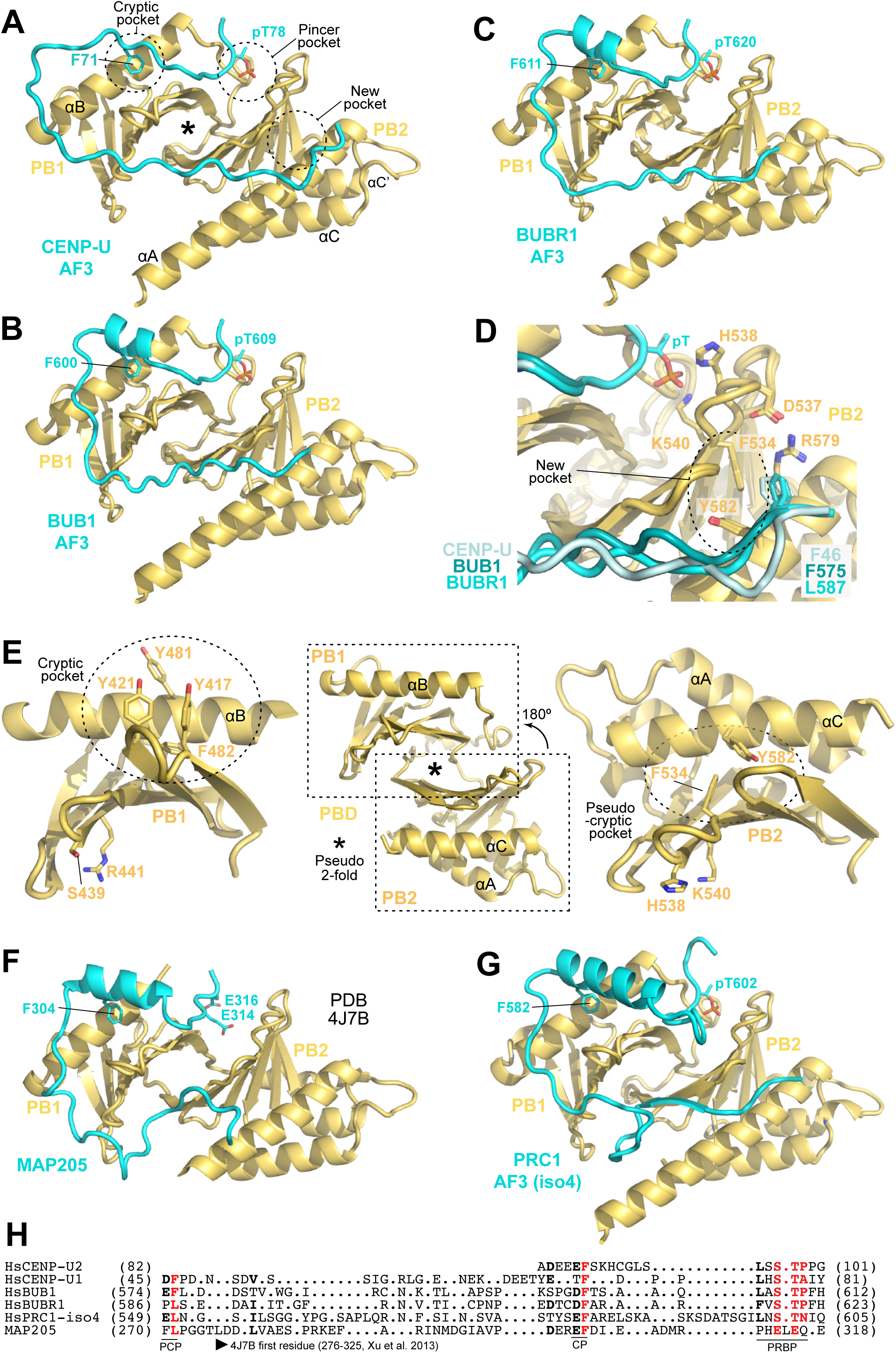
Prediction of PLK1 master docking motifs. (**A**) AlphaFold 3 prediction of a segment of CENP-U. For boundaries of constructs used in this and other predictions in this figure, please refer to Figure S7. For boundaries of displayed segments, refer below to panel H. The asterisk marks the position of a pseudo 2-fold axis discussed in panel E. (**B**) AF3 prediction of a segment of BUB1 bound to the PBD, shown in the same orientation as the model in Panel A, as in successive panels. (**C**) AF3 prediction of a segment of BUBR1 bound to the PBD. (**D**) In this AF prediction, F46 of CENP-U occupies an exposed PB2 pocket. The equivalent indicated residues in predictions with BUB1 and BUBR1 occupy the same pocket. Next to it is a completely conserved salt bridge involving D537 and R579. (**E**) PB1 and PB2 are related by pseudo-2-fold symmetry. We superposed them as explained in the middle panel and show them here in the same orientation on the left (PB1) and right (PB2). This identifies aromatic residues (F534 and Y582) positioned similarly to those in the cryptic pocket, defining the pseudo-cryptic pocket (identified as “New pocket” in panels A-D. It also identifies residues equivalent to the pincer, S439 and R441. Helices are marked to help orientation. (**F**) Crystal structure of the MAP205 peptide of *Drosophila melanogaster* bound to the Zebrafish PBD, extracted from PDB ID 4J7B (Xu et al., 2013). (**G**) AF3 prediction of a segment of PRC1 isoform bound to the PBD. (**H**) ChimeraX (Pettersen et al., 2021) structure-based alignment of the indicated sequences based on the available AF and experimental models. In the underlying annotation: PCP = Pseudo cryptic pocket; CP = Cryptic pocket; PRBP = Phosphoresidue binding pocket. The start residues of MAP205 in the PDB 4J7B file is also indicated.

The previously determined crystal structure of residues 276-325 of MAP205, a PLK1 inhibitor, in complex with a near full-length PLK1 reconstituted from two separate halves (PDB ID 4J7B; (Xu et al., 2013)) strongly reinforced our confidence in these models. In the structure, the MAP205 chain meanders on the surface of the PBD in a manner that is closely related to that observed in the AF3 predictions of the PBD bound to CENP-U, BUB1, and BUBR1. Specifically, F304 of MAP205 positions itself in the cryptic pocket, while two acidic residues (E314 and E316) interact in the pincer pocket, replacing the canonical phosphorylated residue (Figure 4F). The pseudo-cryptic pocket is not occupied in this structure, but a survey of the MAP205 sequence used for crystallization (residues 276-325) showed that it might have been too short. AF3 prediction of a MAP205 fragment with a longer N-terminus (starting at 270) demonstrated the expected occupation of the pseudo-cryptic pocket (Figure S7E).

To put these observations further in context, we referred to PRC1, a known phosphorylation target and binding partner of PLK1 that mediates PLK1 localization to the central spindle in anaphase. We analysed in particular the sequence preceding and including T602 (using Uniprot isoform 4 as sequence reference), a known target of PLK1 phosphorylation (Lim et al., 2024; Neef et al., 2007). Remarkably, AF3 predicted an essentially identical binding mode, also in this case with concomitant occupation of the phosphoresidue-binding site, of the cryptic pocket, and of the pseudo-cryptic pocket (Figure 4G). Collectively, therefore, these observations support the idea that master docking motifs for the PLK1 PBD are much more extended than hitherto appreciated. They can consist of forty or more residues that adopt a U-shape conformation expected to stabilize the PBD very considerably. A tentative structural alignment based on predictions and experimental structures pinpoints the absence of strong sequence constraints, but also identifies similar preferences for the pseudo-cryptic and cryptic pocket, with dipeptides of a negatively charge residue (Asp or Glu) followed by Leu or Phe (Figure 4H). Future structural work should aim to obtain direct experimental evidence for these predictions, but the structure of the PBD:MAP205 complex represents a reassuring precedent. Conversely, comparison with the MAP205 structure makes us suspect that crystal contacts might have forced the N-terminal segment of CENP-U away from its natural course in crystals of the PBD:CENP-U^39-114^ complex.

### Active and inactive PLK1 conformations

An AF prediction of full length human PLK1 shows a closed conformation, with the PBD docked tightly against the kinase domain (Figure 5A). Contacts between the PBD and kinase domain are indirect and mediated by the interdomain-linker (IDL) and the L1 loop. The L2 loop of the PBD, which connects the PB1 and PB2, does not directly contact the kinase domain but is adjacent to the IDL. Because the PBD is known to dampen kinase activity (Chapagai et al., 2023; Jang et al., 2002; Kachaner et al., 2017; Kachaner et al., 2014; Lee and Erikson, 1997; Mundt et al., 1997; Xu et al., 2013), we surmise that the closed structure predicted by AF is that of inactive PLK1, before phosphopeptide-driven activation. This idea is reinforced by a comparison with a previously reported structure of the inhibited PLK1:MAP205 complex, and obtained with two separate halves of PLK1 roughly comprising 1) the KD and part of the IDL, and 2) the PC and PBD domain (Xu et al., 2013). The reciprocal arrangement of the two halves of PLK1 observed in this structure is essentially identical to that of the AF2 prediction of full-length PLK1, implying that MAP205 stabilizes the closed conformation of PLK1.

**Figure 5.**
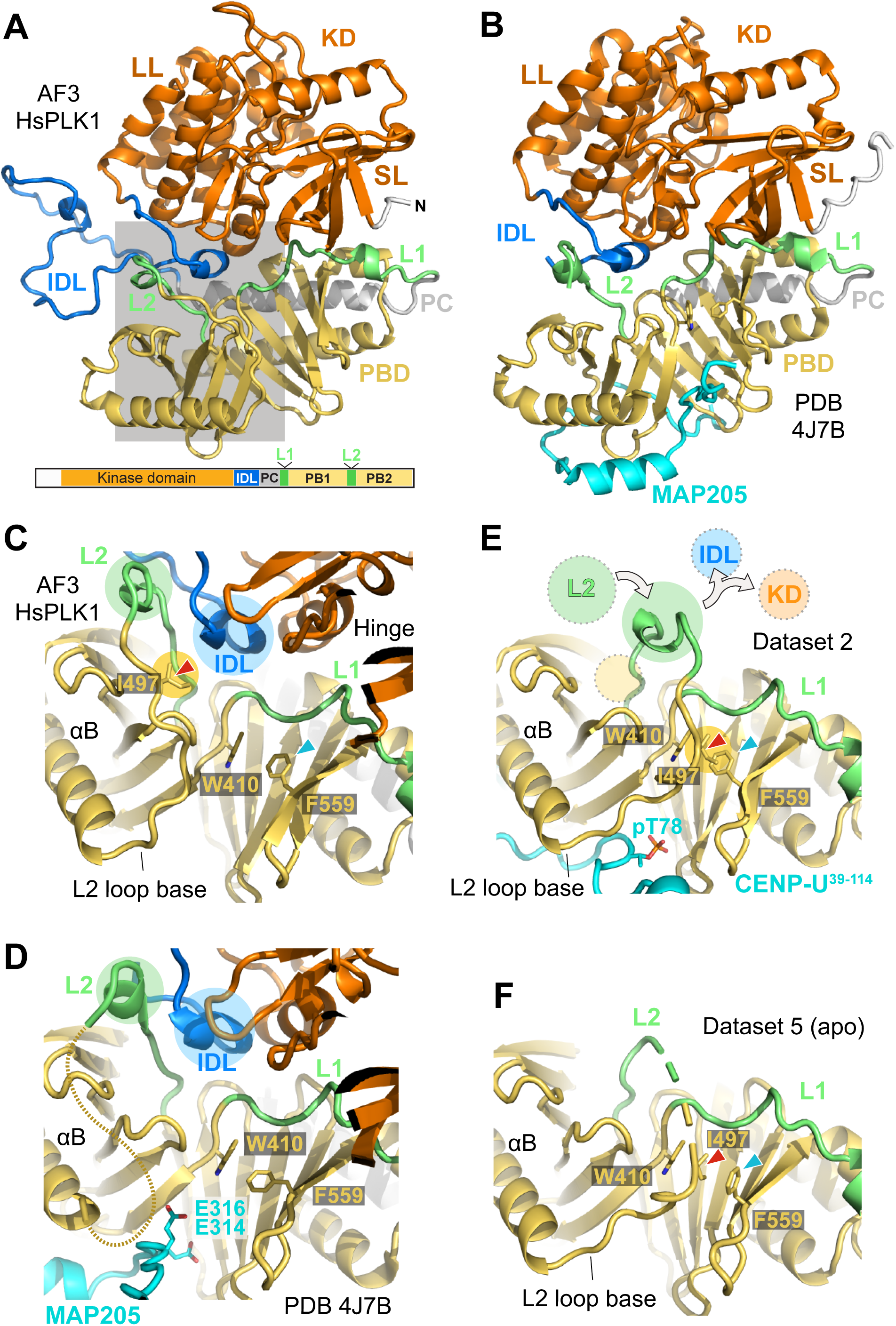
Implications for activation mechanism of PLK1. (**A**) AF modelling of the entire HsPLK1 sequence shows an arrangement that is essentially indistinguishable from that of the PLK1:MAP205 complex. The main structural elements already introduced in Figure 1 are repeated here for reference. The grey area is the focus of closeups shown in panels C-F. SL = Small lobe; LL = Large lobe. (**B**) Cartoon model of the PLK1:MAP205 complex (PDB ID 4J7B). (**C**) Closeup of the AF model of full length HsPLK1, in the grey area of panel A, with rotation to highlight the main elements. The position of the L2 and IDL is indicated with green and blue circles, respectively. The position of the base of the L2 loop is shown. (**D**) PLK1:MAP205 complex. The dashed line represents the expected path of the invisible base of the L2 loop. All elements are in the same position as in the predicted structure of full length PLK1. (**E**) In the structure calculated with dataset 2 (as well as datasets 1, 3, and 4) the L2 loop occupies the position occupied by the IDL in the models displayed in panels C and D. In the context of full length PLK1, the IDL would have to be displaced from its position to permit this conformation of the L2 loop. (**F**) Apo PBD structure calculated from dataset 5. The base of the L2 loop adopts a conformation similar to that observed in the liganded structures exemplified by dataset 2 in panel E. It follows that this conformation is not triggered by peptide binding.

The mechanism of opening and activation of the kinase domain by ligands is a matter of conjecture. None of the AF structures with bound peptides discussed above predicts direct clashes of the extended target peptide with the KD, suggesting an allosteric activation mechanism where binding of phosphorylated target motifs opens the closed arrangement, leading to kinase activation. Target phosphoresidues may change the conformation of the L2 loop, the base of which, at the exit from the αB helix, is juxtaposed to the pincer pocket (Figure 5C-E). In the closed structure predicted by AF, the apex of the L2 loop is stabilized by an interaction with the first segment of the IDL, near the kinase hinge domain (Figure 5C). In the PLK1:MAP205 complex the base of the loop is invisible, but the apex is in essentially the same position as in the AF prediction of full-length PLK1 (Figure 5D).

In our structures of the PBD bound to doubly phosphorylated CENP-U (Table I, datasets 1 and 2), the L2 loop adopts a distinct conformation that is clearly discernible in the available electron density, with weak density only for a few residues centred on Arg500 (Figure 5E and Figure S8A-B). A helpful beacon is the side chain of Ile497, for which there is excellent electron density, and whose position is marked for comparison in the context of the AF model and of the model calculated from dataset 2 (Figure 5C, E). Overall, there is a very significant rotation of the base of the L2 loop, in such a way that in the CENP-U-bound complex the apex of the loop occupies a position overlapping with that adopted by the IDL in the closed conformation, predicting extrusion of the latter and of the associated kinase domain into an unrestrained, active conformation (as exemplified by the arrows in Figure 5E).

To address directly the question whether the conformational change at the base of the L2 loop is related to the presence of a phosphate group on the target motif, we raised crystals of mono-phosphorylated PBD:CENP-U^58-114^ complex (with unphosphorylated T78 and phosphorylated T98) using the co-mixing procedure with both kinases or with only CDK1. For both procedures, crystals grew under largely similar conditions to those of the fully phosphorylated complex obtained by pre-phosphorylation (Table 1, datasets 3 and 4). Structures calculated with datasets 3 and 4 were largely similar to that of the doubly phosphorylated complex (datasets 1 and 2). However, lack of the corresponding electron density indicated unequivocally that T78^CENP-U^ had not been efficiently phosphorylated by PLK1 (Figure 6A-C), while T98^CENP-U^ appeared, as expected, robustly phosphorylated (not shown). A comparison of the organization of the L2 loop in the two new structures indicated that in all cases it adopts conformations that are much more closely related to the open state already observed with datasets 1 and 2 than to the closed, inhibited state (Figure 6D). A cautionary note is that anomalous X-ray scattering (AXRS) detected an unbound free phosphate ion near the position normally occupied by the phosphate group of pT78 in structures from datasets 3 and 4 (indicated by an asterisk in Figure 6B-C). However, we deem it highly implausible that this would be a sufficient trigger to stabilize a particular conformation of the L2 loop. Thus, collectively, this comparison does not appear to support a univocal association between binding of pThr/pSer in the pincer pocket and conformation of the L2 loop. This was further confirmed by the determination of a crystal structure of the apo (unbound) PBD domain (Table 1, dataset 5). While not entirely visible, the L2 loop in the apo-PBD structure adopts a conformation rather similar to that of the liganded PBDs, clustering in an “open” group that is clearly distinct from that expected for the closed assembly (Figure 5F and Figure 6D). Incidentally, the cryptic pocket is occluded in apo PBD, while it is available and occupied by peptide in all eight PBD:CENP-U sub-complexes in our datasets 1 to 4, justifying the name “cryptic” for this pocket (Figure S9A-C).

**Figure 6.**
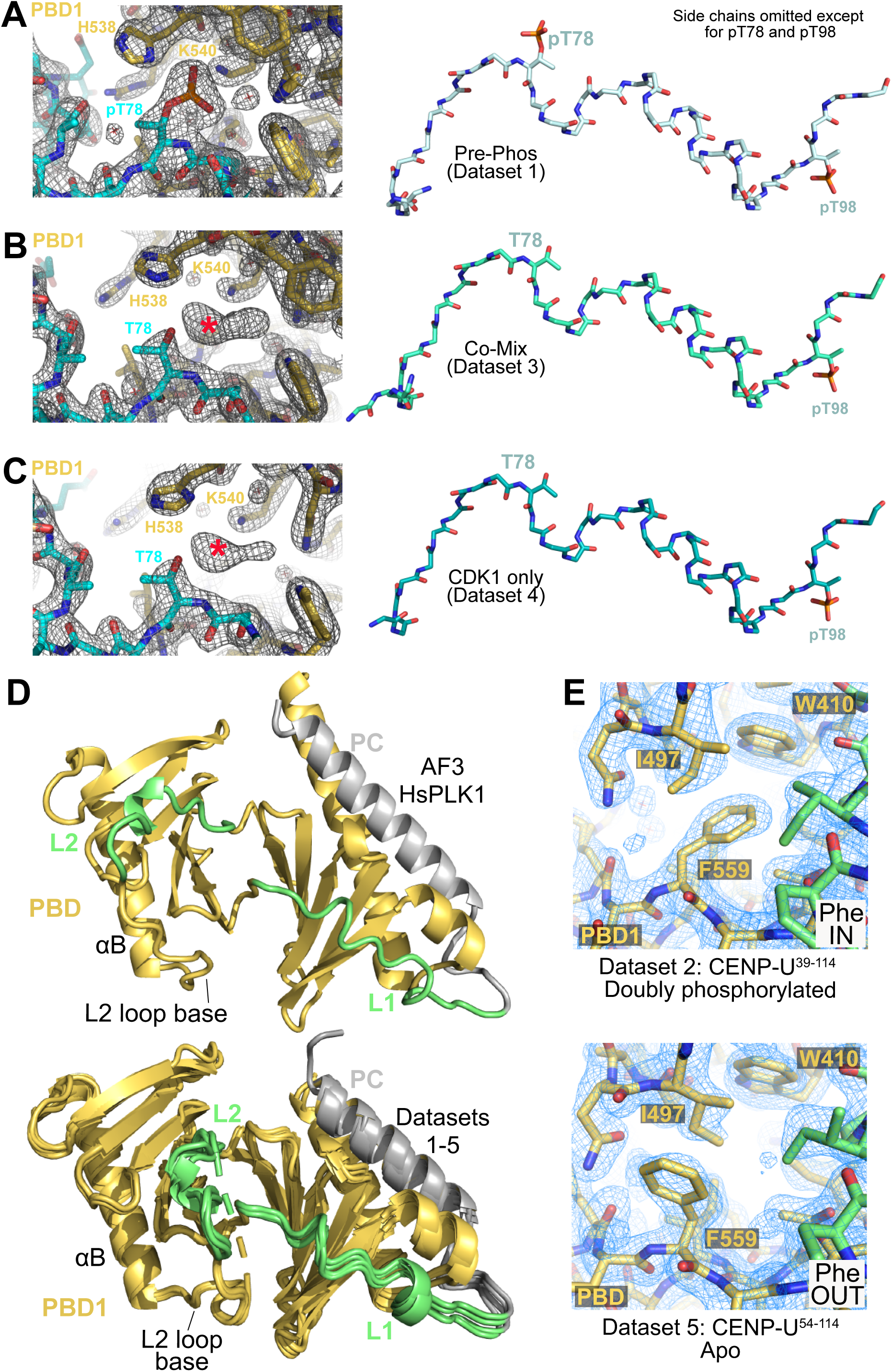
Implications for activation mechanism of PLK1. (**A**-**C**) Comparison of weighted 2F_o_-F_c_ electron densities maps and models calculated with Datasets 1, 3 and 4 (displayed in panels A, B, and C, respectively) demonstrate that despite the absence of phosphate on T78 for datasets 3 and 4, the overall structure of the bound CENP-U peptide (where side chains were omitted to improve clarity) is essentially indistinguishable from that of the doubly phosphorylated peptide. The position of a putative phosphate ion detected by anomalous X-ray scattering (AXRS) is indicated by a red asterisk. Collectively, these structures suggest that structural change introduced by a phosphate on T78 are very small. (**D**) Cartoon model emphasizing the different positions of the L2 loop in the AF model of full-length PLK1 (*top*; other structural elements were removed to improve clarity) and in superposed models calculated from datasets 1 to 5 (*bottom*). (**E**) Phe IN and Phe OUT conformation of F559 documented against the electron density in the indicated datasets. The side chain of Ile497 retains its position in the Phe IN or Phe OUT conformations of Phe559

In the “open” conformation of the L2 loop, the side chain of Ile497 relocates near a pocket formed by Trp410 and Phe559 (Figure 5C-E and Figure 6E). This pocket is the target of a small-molecule allosteric inhibitor of PLK1, Allopole-A, proposed to inhibit ligand binding by dislodging the L2 loop from its position in “open” structures (Park et al., 2023). In isolated PBDs, Ile497 packs against the Trp410 and Phe559 side chains both in peptide-bound and in the apo form (compare Figures 5E-F). The side chain of Phe559 adopts two different conformations in our structures, which we refer to as Phe IN and Phe OUT (Figure 6E). We observe fully Phe IN and Phe OUT conformation in PBD1 of the doubly phosphorylated dataset 2 structure and in the apo dataset 5 structure, respectively (Figure 6E). In other structures (PBD1 and PBD2 in datasets 1, 3, and 4, and PBD2 in dataset 2) the conformations co-exist (Figure S10A-E). Thus, there seems to be no strict correlation between the IN and OUT state of Phe559 and phospho-target binding to the PBD, at least with the isolated PBD. In fact, two other apo structures of the PBD display the Phe IN conformation, contrary to ours (PDB IDs 5NN1 and 1Q4O) (Cheng et al., 2003). Nonetheless, the two conformations of Phe559 are a further indication of the flexibility of the L2 loop.

## Discussion

We confirmed that binding of PLK1 to CENP-U is triggered by initial phosphorylation of T98 and the recruitment of PLK1 to the corresponding medium affinity motif. Subsequent phosphorylation of, and docking on, the otherwise PLK1-kinase-impervious site T78 leads to assembly of a very stable complex. As T78 is not an ideal PLK1 substrate, its resilience to become phosphorylated might be overcome when the kinase is brought in close proximity. T98, on the other hand, is efficiently phosphorylated by CDK1. In a previous study, we proposed dimerization of PLK1 as a putative basis for robust binding to CENP-U that might justify its acting as a master docking site (Singh et al., 2021). Here, we confirm that the PBD can dimerize on CENP-U, while our SEC and SEC-MALS experiments did not detect strong incorporation of two PLK1^FL^ subunits on the doubly phosphorylated CENP-U. We previously reported formation of a PLK1^FL^ dimer on doubly-phosphorylated CENP-U in experiments of analytical ultracentrifugation, and the different outcome of these experiments may be due to differences in buffer and concentration conditions, as well as experimental variability.

Be that as it may, our new observations do not discount dimerization, but do not provide further support for its functional importance, as we find PLK1 binds very tight to a single, dominant motif containing pT78 and dimerization does not increase binding affinity. The affinity measured for the pT78 site is at least 100-fold tighter than the strongest affinities measured so far for phosphorylated motifs *in vitro*, including the pT98-cointaining motif. We argue that the extensive interactions of the CENP-U pT78 motif with the PBD explains this high binding affinity. CENP-U may share this characteristic with a handful of additional master docking sites, including those in BUB1, BUBR1, and PRC1, and future studies will have to address this possibility. Given the rather divergent sequences these motifs display, future work will have to combine further predictions and experimental structure determination to identify crucial binding determinants. With slow dissociation rates, master sites may cause persistent local activation of PLK1 necessary for the phosphorylation of other targets, while providing a hierarchical organization allowing control at only few focal points. We do not expect master sites to be very numerous, however, while secondary sites generated locally may be much more numerous. Indeed, we have demonstrated that CENP-U provides the only high affinity binding site for PLK1 in a reconstituted complex containing the 16-subunit CCAN complex and most of the 10-subunit outer kinetochore complex (Singh et al., 2021).

We and others have recently shown that PLK1 localization to kinetochore in early G1 phase depends entirely on the localization there of M18BP1 (Conti et al., 2024; Parashara et al., 2024). Specifically, after anaphase onset and inactivation of CDK activity, PLK1 becomes recruited to T78 and S93 of M18BP1 through a self-primed mechanism required for deposition of the centromere-specific histone variant CENP-A (Conti et al., 2024; Parashara et al., 2024). Because CENP-U and the rest of the CCAN reside at kinetochores throughout the cell cycle, including early G1 phase, PLK1 would be recruited even in the absence of M18BP1 if CDK1 priming weren’t required for CENP-U localization. Instead, ablation of T78 and S93 phosphorylation on M18BP1 clears PLK1 from anaphase kinetochores. Thus, our current understanding of the mechanisms of PLK1 kinetochore recruitment are consistent with the claim that binding to pT78 primed by PLK1 is only possible after PLK1 docking at the neighbouring pT98 generated by CDK1, a mechanism we define as relay-priming. Why CENP-U utilizes a mechanism of relay priming to generate a master docking site for PLK1 while BUB1 and BUBR1 only utilize direct CDK1 phosphorylation remains unclear. We surmise that it might have to do with the kinetics of dephosphorylation at anaphase, but this will require further analyses. M18BP1 represents another clear example of master docking site, because PLK1, after its recruitment there, can phosphorylate and bind additional proteins involved in the CENP-A deposition pathway. The M18BP1 master site may represent a distinct class with two adjacent phosphorylation sites that bind to a single PBD (Conti et al., 2024). Future work will have to address differences and commonalities between these master docking motifs.

Our work here does not shed new light on how phospho-peptide binding activates PLK1, although it provides an important formal analysis that the contribution of phosphorylation is subtle. The base of the L2 loop is adjacent to the phosphorylated side chain of the target motif, and is therefore the ideal “sensor” of phosphoresidue binding. However, in isolated PBD structures, the conformation of the L2 loop seems to a good degree independent of peptide binding or of its phosphorylation, as shown in Figure 6D. This strongly suggests that the sensitivity to specific features of PBD binding partners must be higher in the context of the full-length PLK1 kinase. MAP205 appears to bind to the PBD in a similarly extensive way as those we have predicted for CENP-U, BUB1, BUBR1, and PRC1, but the outcome is that it stabilizes the closed, less active conformation rather than disrupting it. The crucial difference in MAP205 is the replacement of the S-pS/T-X motif with a Glu-Leu-Glu motif. We suspect that this sequence increases the stability of the closed conformation by locking the base of the L2 loop in a conformation similar to that observed in the AF model of full-length PLK1. In the structure of PLK1 with MAP205, the base of the L2 loop is invisible in the electron density (Xu et al., 2013). This may be a consequence of the deletion of part of the IDL in that structure, as AF predicts stabilizing interactions of the top of the L2 loop with the IDL.

In summary, our observations are consistent with the idea that the L2 loop of PLK1 in the closed, inactive form of PLK1 adopts a more constrained conformation relative to that observed in the structures of isolated PBD domains (with or without binding ligands), which may be considered proxies for the open form of PLK1. The L2 loop conformation in the closed form may be stabilized by reciprocal interactions with the IDL and kinase domain. Its repositioning upon binding of specific targets may provide the energy required to dislodge the IDL and kinase domain, leading to kinase activation. We suspect that master docking sites hold the key to robust activation of PLK1, possibly because by binding to a much larger interface, they elicit a more cooperative and durable stabilization of the L2 loop. These considerations will serve as basis for the design of new studies probing the PLK1 activation mechanism.

## Materials and Methods

### Plasmids

Plasmids encoding 6xHis-MBP-TEV-PLK1^345-603^ (^MBP^PBD), 6xHis-MBP-TEV-CENP-U^58-114^ (^MBP^CENP-U^58-114^), and 6xHis-TEV-PLK1 (PLK1^FL^) for expression in insect cells were available from a previous study and were used precisely as described (Singh et al., 2021). Full length 6xHis-TEV-PLK1 (PLK1^FL^) and CDK1:Cyclin-B:CKS1 (1:1:2) (CCC) were prepared as described in detailed protocols (Huis In ’t Veld et al., 2022; Singh et al., 2021). Plasmids encoding 6xHis-MBP-TEV-CENP-U^84-114^ (^MBP^CENP-U^84-114^), 6xHis-MBP-TEV-CENP-U^58-83^ (^MBP^CENP-U^58-83^) and 6xHis-MBP-TEV-CENP-U^39-114^ (^MBP^CENP-U^39-114^) for bacterial cell expression were constructed by sub-cloning the relative PCR-amplified sequence of codon-optimized CENP-U cDNA in a modified pETDuet-1 (Novagen) in frame with coding sequences for 6xHis and PreScission-substrate–MBP–TEV-substrate. All CENP-U mutants were generated by site-directed mutagenesis directly on these templates.

### Protein expression and purification

^MBP^CENP-U constructs and ^MBP^PBD were expressed in *Escherichia coli* BL21 (DE3) cells. Following transformation with plasmids, the bacteria were cultured in Terrific Broth at 37°C. Protein expression was induced using 0.2 mM IPTG when the optical density reached approximately 0.6 and cells were incubated at 20 °C for approximately 18 hours. Cells expressing PLK1^FL^ and ^MBP^PBD were pelleted and resuspended in Lysis buffer (50 mM HEPES pH 7.5, 500 mM NaCl, 5% (v/v) Glycerol, 10 mM Imidazole, and 1 mM TCEP; or 20 mM Bis-Tris/HCl pH 6.5 in place of 50 mM HEPES pH 7.5 in case of ^MBP^PBD). After supplementation with DNase I (Roche) at 10 μg/mL and either 1 mM PMSF or 160 μg/mL HP plus protease inhibitor mix (Serva), the cell lysates were prepared by fluidization and cleared by centrifugation at 4°C for 45 minutes. The soluble lysate was filtered through a 0.8 µm membrane and applied to a pre-equilibrated, 2× 5-ml HisTALON Cartridge containing pre-packed TALON Superflow Resin (Clontech). The column was washed with at least 10 volumes of the corresponding lysis buffer, and bound proteins were eluted using Lysis buffer supplemented with 300 mM imidazole. Elute fractions were pooled, concentrated and subjected to separation on a Superdex 200 16/60 size exclusion chromatography (SEC) column, pre-equilibrated in SEC buffer 1 (50 mM HEPES pH 7.5, 300 mM NaCl, 5 % (v/v) Glycerol and 1 mM TCEP) or SEC buffer 2 (20 mM Bis-tris/HCl pH 6.5, 200 mM NaCl, 5 % (v/v) glycerol and 1 mM TCEP). Following inspection via 12.5% SDS-PAGE, chosen fractions were concentrated, aliquoted, flash frozen in liquid nitrogen and stored at -80°C. For 6xHis-MBP-TEV-CENP-U variants, bacterial cells were pelleted and resuspended in a Lysis buffer (50 mM HEPES pH 7.5, 500 mM NaCl, 5% (v/v) Glycerol, 10 mM Imidazole, and 1 mM TCEP). Following lysis, cell lysates were initially purified using a HisTALON Cartridge with the His-elution buffer (20 mM Bis-tris/HCl pH 6.5, 200 mM NaCl, 5% (v/v) Glycerol, 200 mM Imidazole, and 1 mM TCEP). Eluates were diluted in 9 volumes of dilution buffer (20 mM Bis-tris/HCl pH 6.5, 5 % (v/v) glycerol and 1 mM TCEP), filtered (0.2 µm membrane), and loaded onto a pre-equilibrated Resource Q anion exchange chromatography column (GE Healthcare) in 20 mM Bis-tris/HCl pH 6.5, 20 mM NaCl, 5 % (v/v) glycerol, 1 mM TCEP. Proteins were then eluted with a linear gradient of 20-1000 mM NaCl and fractions were monitored via 12.5% SDS-PAGE. Selected fractions were combined, concentrated and subjected to further separation using a Superdex 75 16/60 SEC column (GE Healthcare) pre-equilibrated in SEC buffer 2. Following evaluation by SDS-PAGE, chosen fractions were concentrated, divided into aliquots, rapidly frozen in liquid nitrogen, and stored at -80°C.

### In-situ phosphorylation of CENP-U constructs

To functionalize CENP-U variants in their interactions with ^MBP^PBD or PLK1^FL^, we employed 1) a co-mixing protocol where substrate CENP-U variants were incubated with PLK1 constructs immediately before addition of designated active kinase(s) for concurrent phosphorylation and potential complex formation overnight; or 2) a pre-phosphorylation protocol where substrate CENP-U variants were phosphorylated for 15 hours with the designated active kinase(s), and PLK1 constructs were subsequently added to allow potential complex formation. If not otherwise stated, ^MBP^PBD or PLK1^FL^ were added in a molar ratio of 2:1 to the CENP-U variant. All steps were conducted at 4 °C.

### Analytical Size Exclusion Chromatography

SEC analysis was conducted on a Superdex 200 5/150 column. Sample preparation, column equilibration, and elution were performed using SEC buffer (20 mM Bis-tris/HCl pH 6.5, 250 mM NaCl, 5% (v/v) glycerol, and 1 mM TCEP). In case of samples containing PLK1^FL^, the SEC buffer was modified by replacing 20 mM Bis-tris/HCl pH6.5 with 20 mM HEPES pH7.5. Before loading on the column, a fraction of the sample was held as input for SDS-PAGE with Pro-Q™ Diamond and Coomassie blue staining. For direct interaction analysis between binding-functionalized CENP-U variants and PLK1^FL^, sample inputs were verified further by Western blotting using antibodies specific to the phosphorylated Thr78^CENP-U^ (anti−PBIP-1(phosphor-T78) rabbit polyclonal; Abcam, Cambridge, UK; 1:1000 diluted for use) and phosphorylated Thr210^PLK1^ (anti-PLK1-phospho-T210, mouse monoclonal; Biolegend #629801, 1:1000 diluted for use). All samples were eluted under isocratic conditions at 4°C with a flow rate of 0.15 ml/min. Protein elution was monitored by UV detection at a wavelength of 280 nm and 100 µl fractions were collected that were subsequently analysed by SDS-PAGE with TCE (2,2,2-Trichlorethanol) or Coomassie blue staining in case of samples containing PLK1^FL^. Stained gels were digitalized using a BioRAD chemiDoc MP Imaging System (BioRAD).

### Multi Angle Light Scattering

SEC-MALS analyses were performed using a 1260 Infinity II SEC system coupled with a Dawn Heleos-II System and an Optilab T-rEX Refractive Index (RI) detector (Wyatt). A Superdex 200 10/300 column (GE Healthcare) was used for the SEC elution and pre-equilibrated with Glycerol-free SEC buffer containing 20 mM Bis-tris/HCl pH 6.5, 250 mM NaCl and 1 mM TCEP, or in case of samples involving PLK1^FL^, 20 mM HEPES pH7.5, 300 mM NaCl and 1 mM TCEP. The column was cooled to 12 °C using a Shimadzu column cooler. Samples were prepared with a total protein concentration higher than 2 mg/mL and of which 60 μL were loaded for the analysis. The theoretical dn/dc for each sample was calculated using SedFit. The analysis was done by Astra 7.3.2.21, using BSA for peak normalization. The molar mass for each peak was calculated using the theoretical dn/dc of the proteins and the RI signal. The data were fitted with the Zimm equation.

### Preparation of phosphorylated CENP-U variants

CENP-U variants (in 100 μM) were phosphorylated by incubation with the indicated kinase overnight at 4°C in phosphorylation buffer (with the same composition as SEC buffer 2) supplemented with MgCl_2_ (10 mM) and ATP (2.0 mM). CCC was used at a 1/30^th^ of the concentration of substrate. The corresponding conversion rate of this Thr98 phosphorylation method was determined by mass spectrometry to exceed 96 % (LR and PJ, unpublished observations). Thr78 was phosphorylated using activated 6xHis-TEV-PLK1 (PLK1^FL^) activated *in vitro* with Aurora-A/Bora complex at a molar ratio of 200:10:1 (PLK1:Aurora-A:bora) as recently described (Lim et al., 2024). The corresponding conversion rate of this Thr78 phosphorylation method was determined by mass spectrometry to be around 90 % (LR and PJ, unpublished observations). After incubation the reaction mixture was then concentrated and purified using Superdex 75 16/60 SEC column (GE Healthcare) pre-equilibrated in the SEC buffer. Fractions were monitored by SDS-PAGE and Pro-Q™ Diamond staining (Invitrogen, according to the manufacturer instruction), those containing the phosphorylated protein were concentrated, flash-frozen in liquid nitrogen and stored at -80°C.

### Preparation of CENP-U:PBD complexes for crystallographic analysis

In preparation for crystallization, we first assembled pre-phosphorylated the relevant ^MBP^CENP-U fragments on ∼15 mg scale (at 40 μM reaction concentration) and incubated them with CDK1 and PLK1 for 15 hours. We then incubated the phosphorylated CENP-U with 2 equivalents of ^MBP^PBD for ∼3 hr in assembly buffer (20 mM Bis-tris/HCl pH 6.5, 300 mM NaCl, 5 % (v/v) glycerol, 1 mM TCEP), further concentrated the mix, and separated it on a Superdex 200 Increase 10/300 SEC column (Cytiva) pre-equilibrated in the assembly buffer. Fractions containing assembled complex were collected and concentrated to roughly the previous volume of the assembly mixture. The 6xHis-MBP-tagged TEV Protease (made in house) was added at 1/20 dilution to substrates and cleavage was allowed to run for 15 hours at 4°C. The mixture was diluted with dilution buffer (20 mM Bis-tris/HCl pH 6.5, 5 % (v/v) glycerol, 1 mM TCEP) to achieve a NaCl concentration of 50 mM and loaded onto a tandem 5-mL HisTALON Cartridge (Clontech) and Resource Q anion exchange chromatography column (GE Healthcare) pre-equilibrated in 20 mM Bis-tris/HCl pH 6.5, 50 mM NaCl, 5 % (v/v) glycerol, 1 mM TCEP. The complex was eluted using a linear gradient ranging from 50 to 1000 mM NaCl. Following SDS-PAGE analysis with Pro-Q™ Diamond and Coomassie blue staining, fractions containing PBD and CENP-U (the latter only visible upon phosphoprotein staining) were pooled and concentrated for a final SEC purification and buffer exchange on a Superdex 75 10/300 SEC column (GE Healthcare) pre-equilibrated in Xtal-SEC buffer (10 mM Bis-tris/HCl pH 6.5, 100 mM NaCl, 1 mM TCEP). After elution, the fractions were examined again by SDS-PAGE with Pro-Q™ Diamond phosphoprotein staining and Coomassie blue staining. The purified complex was concentrated to ∼8 mg/mL and used freshly for crystallization screen. The same protocol was adapted for the co-mixing procedure.

### Crystallization and structure determination

Protein complexes used for crystallization were freshly purified. Crystallization was conducted using the sitting-drop vapor diffusion method in TTP-IQ format 96-well plates (SPT LabTech) at 20 °C. The screening-plate setup was performed utilizing a mosquito HTS robot (SPT Labtech) at 20 °C, wherein protein sample was mixed with reservoir crystallization solution in a volume ratio of 1:1 or 2:1. Crystals of the doubly phosphorylated complex containing CENP-U^58-114^ appeared within 24 hrs and kept growing for 4 days in a solution containing 0.1M CHES pH 9.5, 0.2 M NaCl, 10% PEG8000. The same solution containing 15% (v/v) PEG400 was then used as cryoprotectant for a brief soak-in of crystals before mounting onto a cryo-loop and promptly flash freezing in liquid nitrogen. Crystals of the monophosphorylated complex emerged grew rapidly (less than 1 day) in 0.08 M NaH2(PO4) pH 6.2, 0.02 M Na3-citrate and 18% PEG2000. 10% (v/v) PEG400 was used as cryoprotectant. Crystals of doubly phosphorylated complex containing CENP-U^39-114^ appeared within 6 hrs and grew for 2 days in a solution containing 0.2 M tri-Lithium citrate, 20% PEG3350. 10% (v/v) glycerol was used as cryoprotectant. Collection of native synchrotron X-ray diffraction data for and anomalous X-ray scattering assay (AXRS) for phosphorus atom determination (Cianci et al., 2005) were conducted at beamline PXII X10SA at the Suisse Light Source in Villigen (Switzerland) or at the ID23-2 microfocus beamline at European Synchrotron Radiation Facility (ESRF) in Grenoble (France). Diffraction intensities were processed using XDS and scaled with XSCALE (Kabsch, 2010). Molecular replacement (MR) using the reported crystal structure 1Q4K (PDB ID) as the search template (Cheng et al., 2003) was done with MR Phaser (McCoy et al., 2007), followed by map refinement with Phenix.refine (Adams et al., 2010). Manual model building was performed with Coot (Emsley et al., 2010). The structure of the doubly phosphorylated complex was subsequently utilized for phasing all other acquired datasets. All crystallographic statistics are summarized in Table 1. X-ray structures were analysed and corresponding presentation images were created with Pymol 2.60 software (Schrödinger LLC, New York, NY).

### Biotin labelling and Bio-layer Interferometry (BLI)

Biotin-Streptavidin (SA) chemistry was used for immobilization of proteins onto the tip surface of an OctetRed 384 sensor from Sartorius. A DNA sequence encoding a PLPETG peptide was inserted in frame at the 3’ end of the coding region for chosen immobilized ligand proteins (Table 2), and the same protocols were used for purification. The C-terminal peptide was conjugated with Gly-Gly-Gly-Gly-Biotin (Thermo Fisher Scientific) for Biotin labelling using calcium-independent 7+ sortase A (Jeong et al., 2017). Reaction was conducted for 15 hours at 4°C in a molar loading ratio of 1:10:100 (sortase:protein:label) in SEC buffer of the specific ligand protein. The biotinylated ligand protein was separated from the catalytic enzyme and the excess label in the mixture by SEC in SEC buffer 2 and subsequently flash-frozen in liquid nitrogen and stored at -80°C. To minimize buffer mismatch, all proteins, including the ligand protein and analyte proteins for titration, were subjected to buffer exchange through dialysis in BLI dialysis buffer (20 mM HEPES pH 7.5, 2.5 % (v/v) Glycerol, 300 mM NaCl, 0.1 mM TCEP). Concentrations of the dialyzed proteins were re-measured immediately before performing the BLI assay. The initial highest concentration of either the ligand or any analyte for BLI assay was achieved by diluting the dialyzed protein with BLI dialysis buffer and 5x BLI additive buffer (20 mM HEPES pH 7.5, 2.5 % (v/v) Glycerol, 300 mM NaCl, 0.1 mM TCEP, 1.5 g/L BSA, 0.25% (v/v) Tween20), ensuring a final buffer containing 0.3 g/L BSA and 0.05% (v/v) Tween20. Other dilutions were performed using the BLI working buffer (20 mM HEPES pH 7.5, 2.5 % (v/v) Glycerol, 300 mM NaCl, 0.1 mM TCEP, 0.3 g/L BSA, 0.05% (v/v) Tween20) in 2- or 3-fold serial dilutions. BLI data were recorded with the Octet BLI Discover 13.0 software. The temperature was maintained at 25°C throughout the experiments and plates/wells were agitated at 1000 r.p.m. when sensors were immersed. The protein was immobilized on Streptavidin SA sensors (Sartorius). The data were analysed using Octet Analysis 13.0. Kinetic signals were initially fitted using a local method. In general, measurements with a local fitting R^2^ value above 0.95 and an RSS value below 0.01 were deemed of high quality and selected for subsequent global fitting. The final affinity constant (K_D_) was calculated as the ratio of k_off_ and k_on_ values given by the global fitting.

### Isothermal Titration Calorimetry

ITC experiments were performed with a PEAQ-ITC calorimeter (Malvern Panalytical). Protein samples were thoroughly dialyzed or subjected to gel filtration at 4 °C against the ITC buffer, containing Hepes (pH 7.5, 20 mM), NaCl (300 mM), TCEP (1 mM) and Glycerol (1%). Following concentration adjustment, the sample cell was loaded with the specified phosphorylated CENP-U construct and titrated by ^MBP^PBD in the injection syringe at 25 °C. The stirrer speed was set to 750 rpm. As a control, ^MBP^PBD at similar concentrations was titrated into ITC buffer. The data was analyzed using Microcal PEAQ-ITC 1.41 (Malvern Panalytical). Control measurements were subtracted from the experimental data using line fitting of the heat peaks. The dissociation constant was determined using Microcal PEAQ-ITC 1.41 employing the one set of sites binding model (Wiseman et al., 1989).

### Mass spectrometry (MS)

The phosphorylation status of CENP-U variants resulting from different *in-vitro* phosphorylation treatments with kinase CDK1 and/or kinase PLK1 were assessed using liquid chromatography coupled mass spectrometry (LC-MS). Controls were included for all samples by omitting the kinase(s). Samples were treated with 1 mM Dithiothreitol and alkylated with 5.5 mM chloroacetamide, followed by sequential digestions using LysC and Trypsin. After termination with TFA (0.15% (v/v)) the resulting cut peptides was purified and enriched using StageTips as previously described (Rappsilber et al., 2007). For MS analysis, 200 ng (CENP-U based) of the obtained cut-peptide mixtures, after passage through a desalting cartridge with 0.1% formic acid (aq), were separated on a Pepmap C18 nanoHPLC column (U3000 nanoHPLC system, Thermo Fisher Scientific) by performing a gradient elution using 5-30% acetonitrile with 0.1% formic acid at a flow rate of 300 nl/min. Eluted liquid was directly sprayed into an Orbitrap type mass spectrometer via a nano-electrospray source in a Q Exactive (Thermo Fisher Scientific). A data-dependent mode was applied for one survey scan, followed by up to 10 MS/MS scans (Olsen et al., 2007). To identify phospho-sites, the resulting raw files were processed with MaxQuant (version 2.2.0.0) searching for CENP-U sequences with acetylation (N-term), oxidation (M) and phosphorylation (STY) as variable modifications and carbamidomethylation (C) as fixed modification (Cox and Mann, 2008). A false discovery rate cut-off of 1% was applied at the peptide and protein levels and as well on the phosphorylation site table (Cox and Mann, 2008). Relative quantification of the phosphorylated peptides of interest, such as DEETYETFDPPLHSp**T**AIYADEEEFSK for Thr78^CENP-U^ and HCGLSLSSp**T**PPGK for Thr98^CENP-U^, and of the unphosphorylated counterparts were performed using Skyline (MacLean et al., 2010; Schilling et al., 2012). The msms.txt table of the MaxQuant search was used to build-up the library and all peak quantifications were checked manually in Skyline. If necessary peak areas were re-integrated manually. Peaks with highest localization probability of phosphorylation of positions Thr78 or Thr98, respectively were extracted. MS intensities of the peptide with Thr78 phosphorylated and of the unphosphorylated counterpart were used for the analysis of PLK1-mediated phosphorylation, and so were those of Thr98 used for the analysis of CDK1-mediated phosphorylation. The phosphorylation conversion rates were determined based on the Thr-unphosphorylated peptide in samples using the formula, 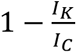, where IK is the MS intensity of the unphosphorylated peptide from a kinase-treated sample, while IC is that from the control sample.

### AlphaFold modelling

Alphafold multimer 2.3.1 (AF2) (Jumper et al., 2021) and Alphafold 3.0.0 (AF3) (Abramson et al., 2024) were used for all structure predictions. Post-translational modifications (i.e. phosphothreonine and phosphoserine) and ligands (ATP) were defined using CCD codes TPO, SEP and ATP. For AF2 predictions, ten models per run were ranked according to the TM score and analyzed, for AF3 five models. PAE plots were generated using a heavily modified version of “Alphafold-analysis” (https://github.com/grandrea/Alphafold-analysis) and used together with the pLDDT scores to assess the quality of the models.

## Author contributions

Conceptualization LR, AEV, IV, AM

Formal Analysis LR, IV

Funding acquisition AM

Investigation AEV, CK, FM, IV, LR, RG, MP, PJ, PG, SW

Project Administration LR, AM

Resources AEV, CK, SW

Supervision AM, LR

Visualization AM, LR, IV

Writing original draft AM, LR

Writing - review & editing All authors

## Acknowledgements

We would like to thank the technical staff of the Swiss Light Source at the Paul Scherrer Institut, Villigen, Switzerland for their support. We acknowledge the European Synchrotron Radiation Facility (ESRF) for provision of synchrotron radiation facilities under proposal number mx2491 and we would like to thank A. Gautam for assistance and support in using beamline ID23-2. Atomic coordinates obtained from the five datasets have been deposited with the protein data bank under accession code 9FJH, 9FJG, 9FJI, 9FJJ, 9QMO. A.M. acknowledges funding from the Max Planck Society, the European Research Council (ERC) Synergy Grant 951430 (BIOMECANET), the DFG’s Collaborative Research Centre 1430 "Molecular Mechanisms of Cell State Transitions", and the CANTAR network under the Netzwerke-NRW program.

## Supplementary Figure Legends

**Figure S1.**
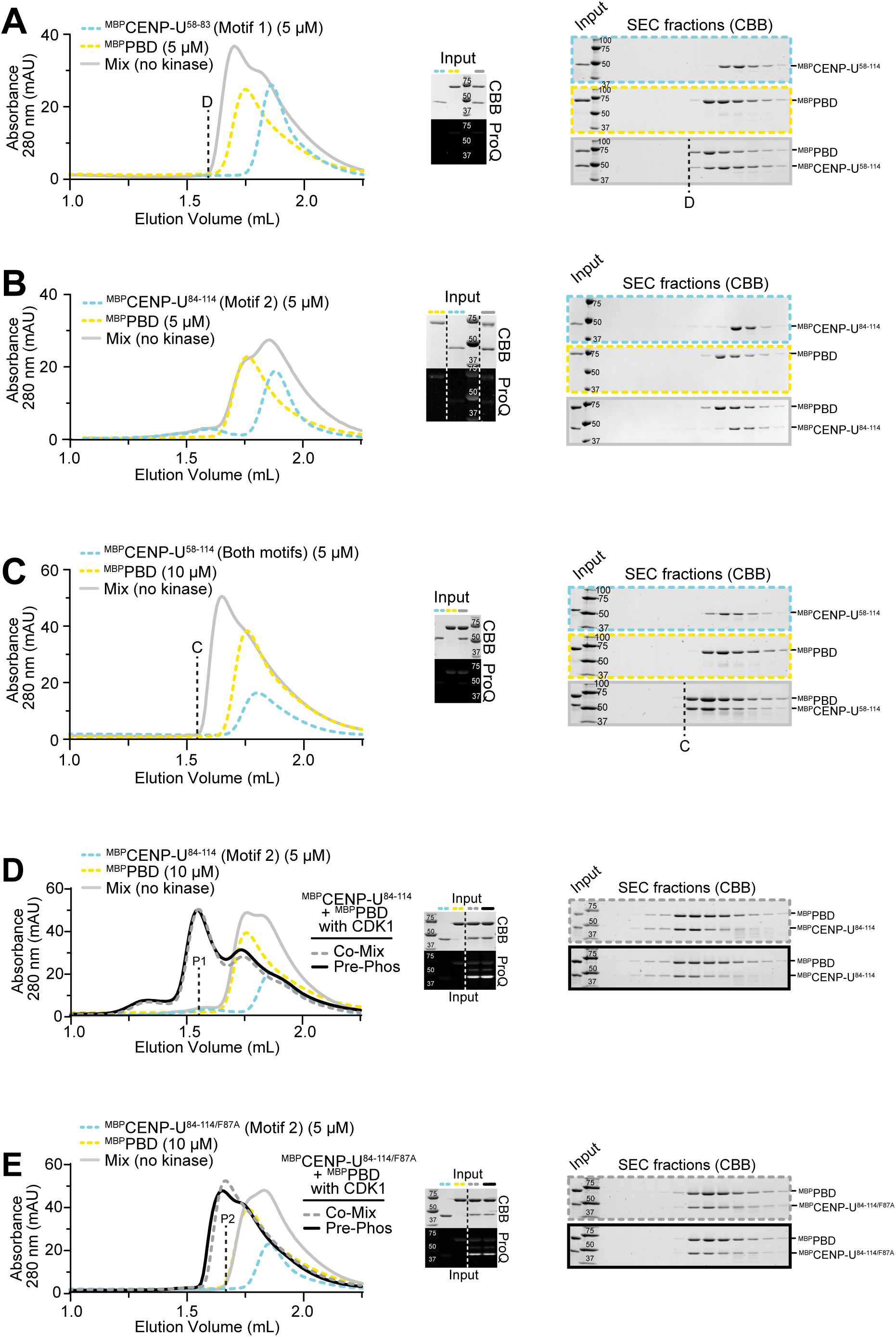
**(Associated with Figure 1)** (**A**) Analytical SEC profiles coupled with corresponding SDS-PAGE and Pro-Q™ analyses for the determination (along with panel B-C) that PBD has residual affinity for unphosphorylated Motif 1, but not Motif 2. D indicates the elution front of the weak complex of PBD with Motif 1. (**B**) No co-elution front of PBD with Motif 2 was observed. (**C**) C indicates the elution front for the weak complex of PBD to the unphosphorylated ^MBP^CENP-U^58-114^ (covering both Motif 1 and 2). (**D**) Demonstration (along with panel E) that the hydrophobic residue (F87^CENP-U^) is required for stable binding between Motif 2 and PBD. P1 indicates the elution peak of the stable complex of the PBD with the phosphorylated Motif 2. (**E**) P2 indicates the elution peak of the weak complex of PBD with phosphorylated Motif 2 carrying the F87 ^CENP-U^ mutation. Both *in-situ* phosphorylation methods (see Methods) were applied for protein phosphorylation displayed in panels D and E.

**Figure S2.**
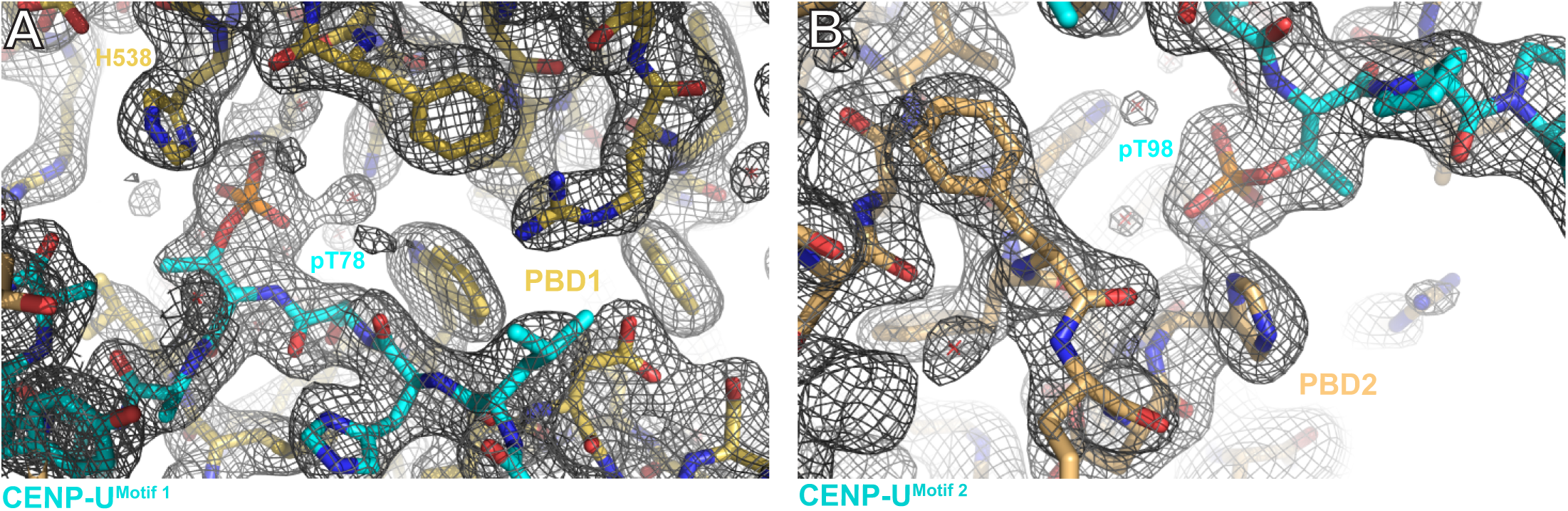
**(Associated with Figure 2)** (**A**) Weighted 2F_o_-F_c_ electron density map calculated from dataset 1 of CENP-U Motif 1 bound to PBD1, centred at pT78. (**B**) Weighted 2F_o_-F_c_ electron density map calculated from dataset 1 of CENP-U Motif 1 bound to PBD1, centred at pT98.

**Figure S3.**
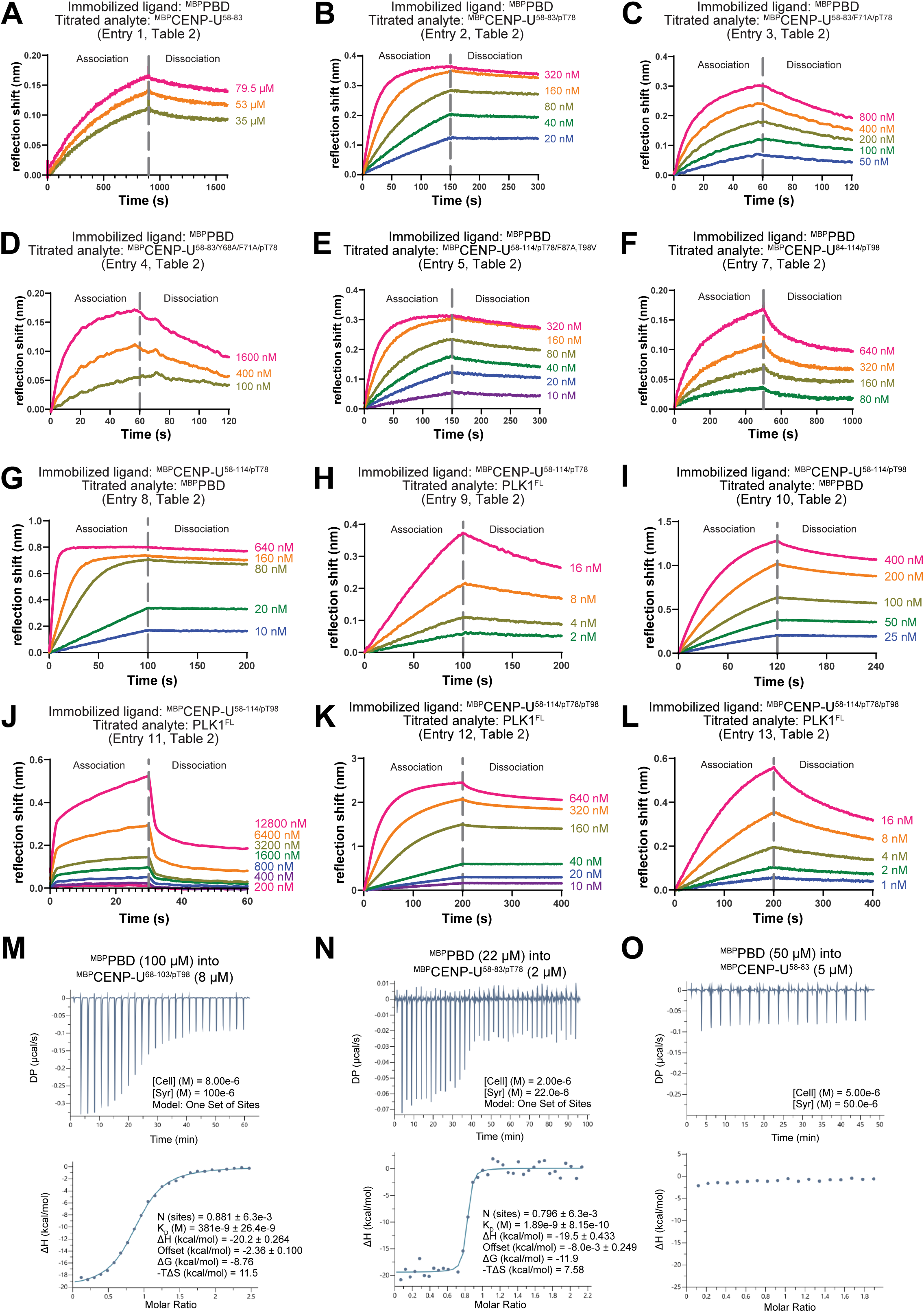
**(Associated with Table 2)** (**A**-**L**) Sensorgrams of the indicated BLI experiments. (**M**-**O**) Thermograms of ITC measurements. ITC titrations of (**M**) ^MBP^CENP-U^68-103/pT98^, (**N**) ^MBP^CENP-U^58-83/pT78^, and (**O**) and ^MBP^CENP-U^58-83^ with ^MBP^PBD. With the applied protein concentrations and buffer conditions in these ITC assays, no notable heat release was observed in the titration of the T78-site unphosphorylated ^MBP^CENP-U^58-83^ with ^MBP^PBD, supporting single-site fitting model of the data calculation in panel M.

**Figure S4.**
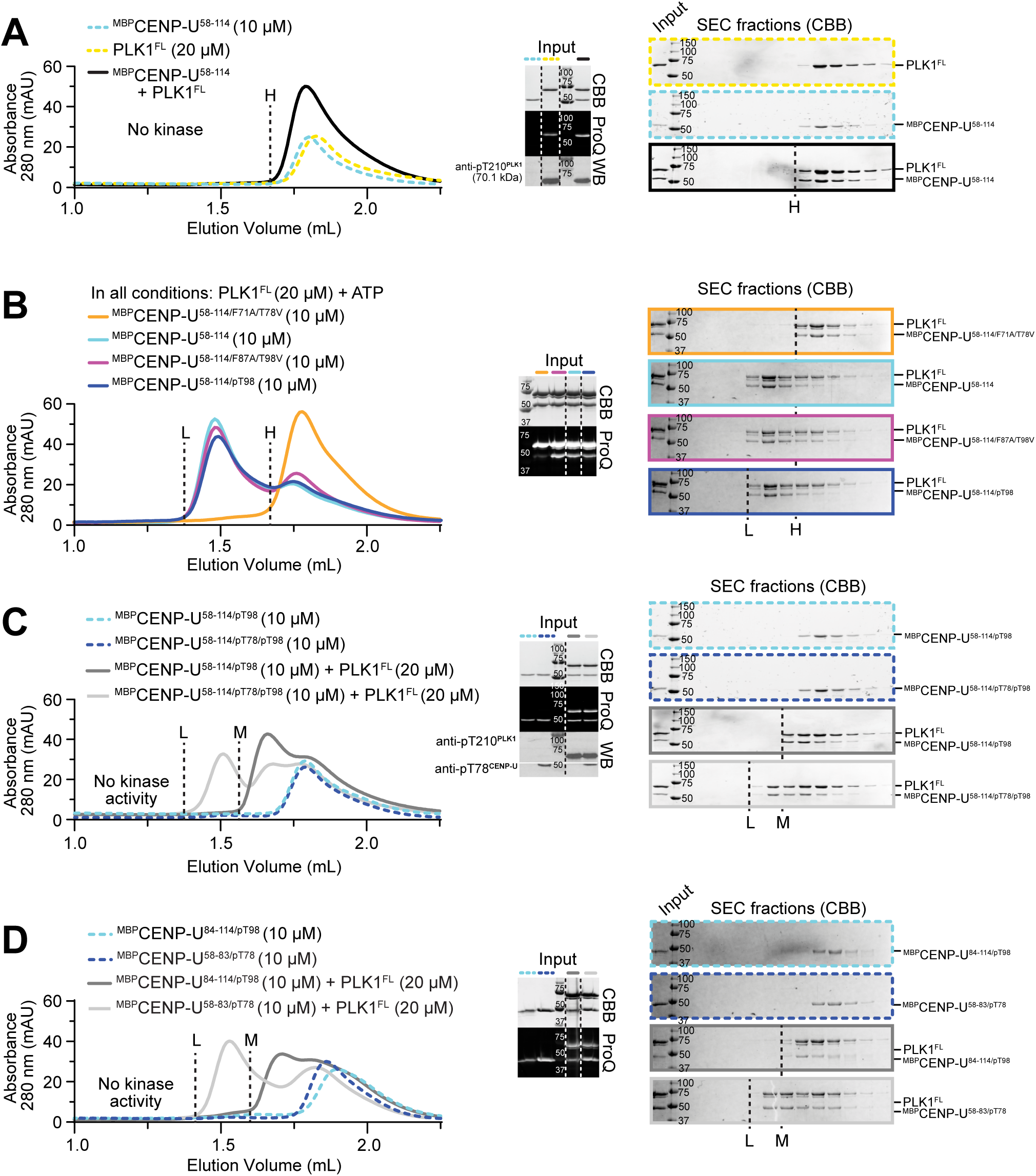
**(Associated with Table 2)** (**A**) Analytical SEC profiles and corresponding SDS-PAGE and Pro-Q™ analyses demonstrating interactions of CENP-U pT78-Motif 1 and pT98-Motif 2 with PLK1^FL^. Elution front H suggests no obvious affinity of PLK1^FL^ to ^MBP^CENP-U^58-114^ (covering both Motif 1 and 2). The PLK1^FL^ used in all panels A-D are active, as shown by phosphorylation of Thr210 demonstrated by Western blotting. (**B**) PLK1^FL^, at 20 μM concentration, directly phosphorylates T78 of CENP-U (10 μM) upon addition of ATP, as shown by Pro-Q™ analyses. The protein concentrations are consistent in all panels. As indicated by elution front L, the CENP-U peptide and CENP-U mutated at Motif 2 (^MBP^CENP-U^58-114/F87A/T98V^) were phosphorylated and formed heterodimer with PLK1^FL^. CENP-U mutated at motif 1 (^MBP^CENP-U^58-114/F71A/T78V^) can barely be phosphorylated (Pro-Q™) and showed no interaction with PLK1^FL^ (elution front H). Notably, T98-prephosphorylated CENP-U (^MBP^CENP-U^58-114/pT98^), subject to the same PLK1 phosphorylation treatment to phosphorylated T78, bound only one PLK1^FL^ (elution front L), confirming that dimerization of PLK1^FL^ on CENP-U is not as stable as dimerization of the PBD. (**C**) In absence of ATP-Mg^2+^, doubly-phosphorylated CENP-U (^MBP^CENP-U^58-114/pT78/pT98^, validated by Western blotting) also bound a single PLK1^FL^ (elution front L). Notably, T98-prephosphorylated CENP-U, without phosphorylation at T78, revealed a weaker interaction with PLK1^FL^ (elution front M). (**D**) In contrast with the T98-prephosphorylated CENP-U (elution front M, also shown in panel C), T78-prephosphorylated CENP-U showed strong binding with PLK1^FL^ (elution front L).

**Figure S5.**
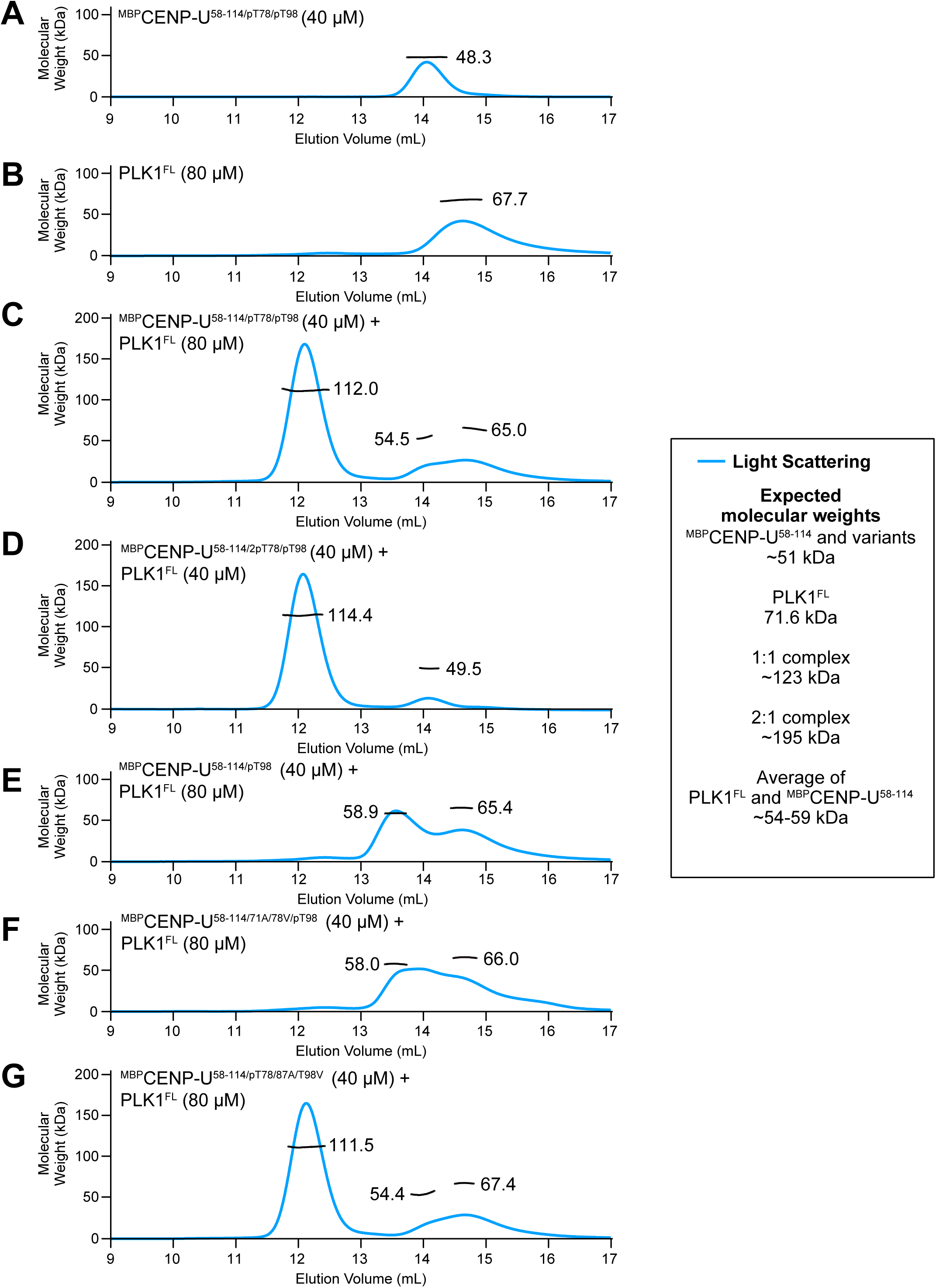
**(Associated with Table 2)** (**A**) SEC-profiles and average size-based molecular weight (MW) in SEC-MALS measurement of the pre-prepared (^MBP^CENP-U^58-114/pT78/pT98^, see also Figure S4C). The calculated MW for the monomer is 51.2 kDa (see legend). (**B**) SEC-MALS analysis of the PLK1^FL^ monomer. (**C**) SEC-MALS analysis of the doubly-phosphorylated CENP-U (panel A) with PLK1^FL^ (panel B) detected a 1:1 complex, despite a 1:2 molar mixing ratio. (**D**) SEC-MALS of the same sample generated at a mixing ratio of 1:1 gave a similar result. (**E**) SEC-MALS of the T98-prephosphorylated CENP-U (^MBP^CENP-U^58-114/pT98^) and PLK1^FL^ delivered a MW suggesting a weak, consistent with results in Figure S4C-D. (**F**) A result similar to that obtained in panel E was obtained when Motif 1 was mutated and T98-prephosphorylated with CDK1 (^MBP^CENP-U^58-114/F71A/T78V/pT98^). (**G**) In contrast to the results in panel F, SEC-MALS revealed strong binding of PLK1^FL^ mixed 1:1 with CENP-U prephosphorylated on T78 and mutated on Motif 2 (^MBP^CENP-U^58-114/pT78/F87A/T98V^).

**Figure S6.**
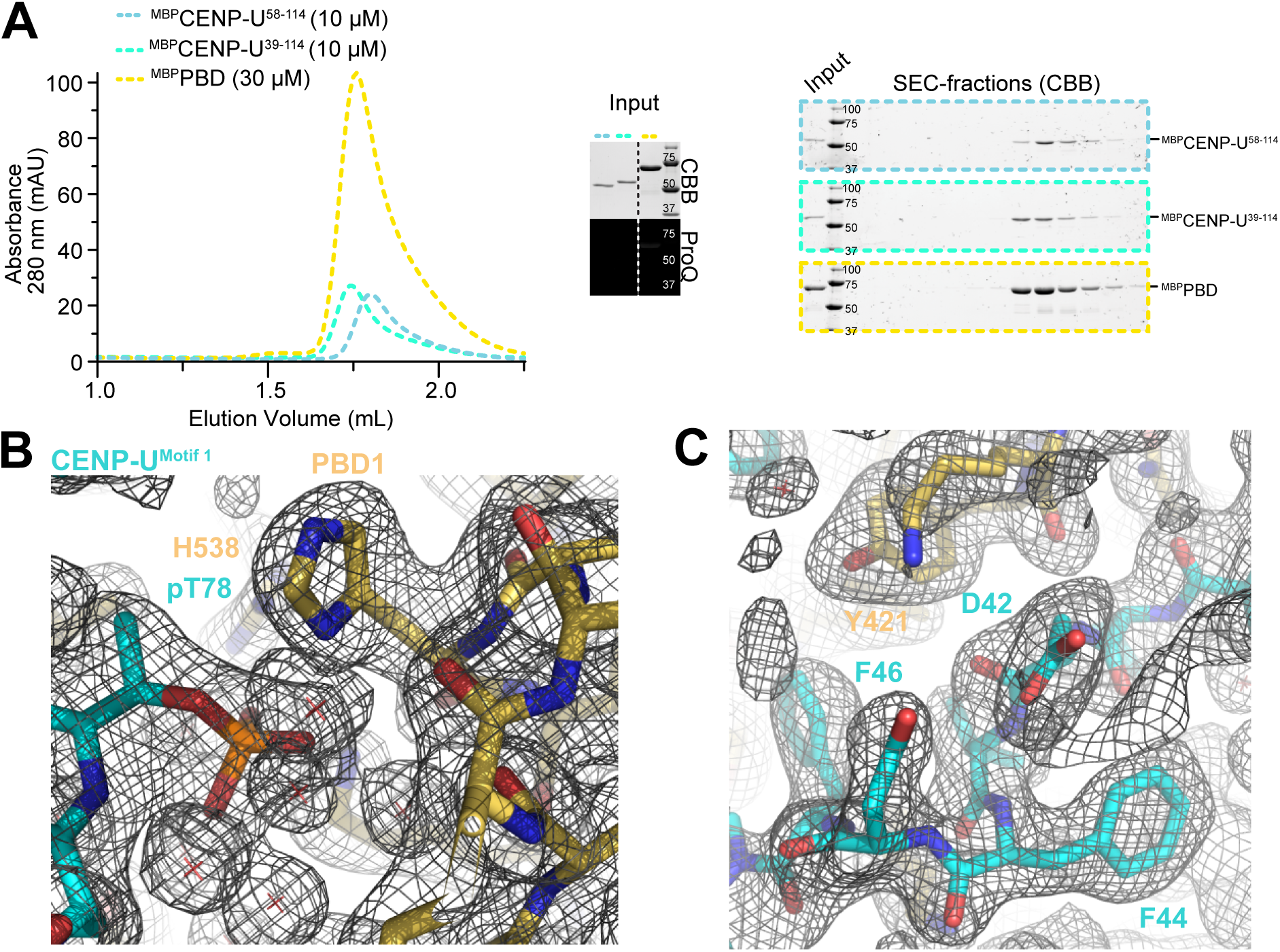
**(Associated with Figure 3)** (**A**) Analytical SEC profiles coupled with corresponding SDS-PAGE and Pro-Q™ analyses of control protein samples used for the experiments in Figure 3B. (**B**) Weighted 2F_o_-F_c_ electron density maps of interacting spaces in the crystallized complex of N-terminal extended ^MBP^CENP-U^39-114^ in compex with two PBDs (Table 1, dataset 2; Figure 3C). T78^CENP-U^ phosphorylation was clearly detected. The phosphorylated side chain interacts with the canonical pincer residues of the PBD. (**C**) The weighted 2F_o_-F_c_ electron density map of the N-terminal extended region (residues 39-47) of CENP-U demonstrates extensive contacts with the PBD.

**Figure S7.**
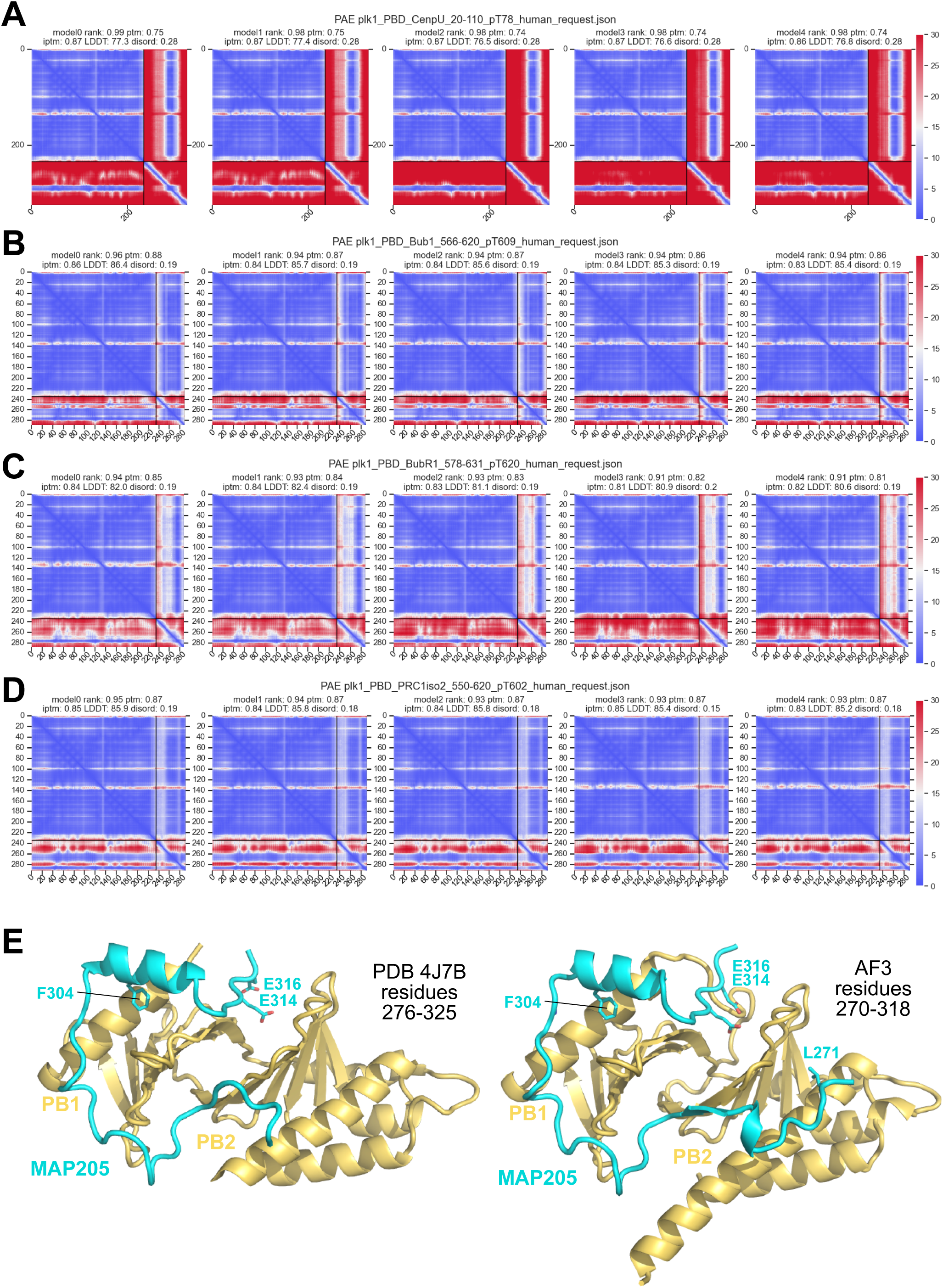
**(Associated with Figure 4)** (**A**-**D**) AF LDDT scores for the indicated predictions displayed in Figure 4. (**E**) Comparison of crystal structure of the PBD-MAP205 complex and of the AF3 prediction with a peptide containing an N-terminal extension. AF3 predicts that the longer peptide occupies the pseudo-cryptic pocket with L271.

**Figure S8.**
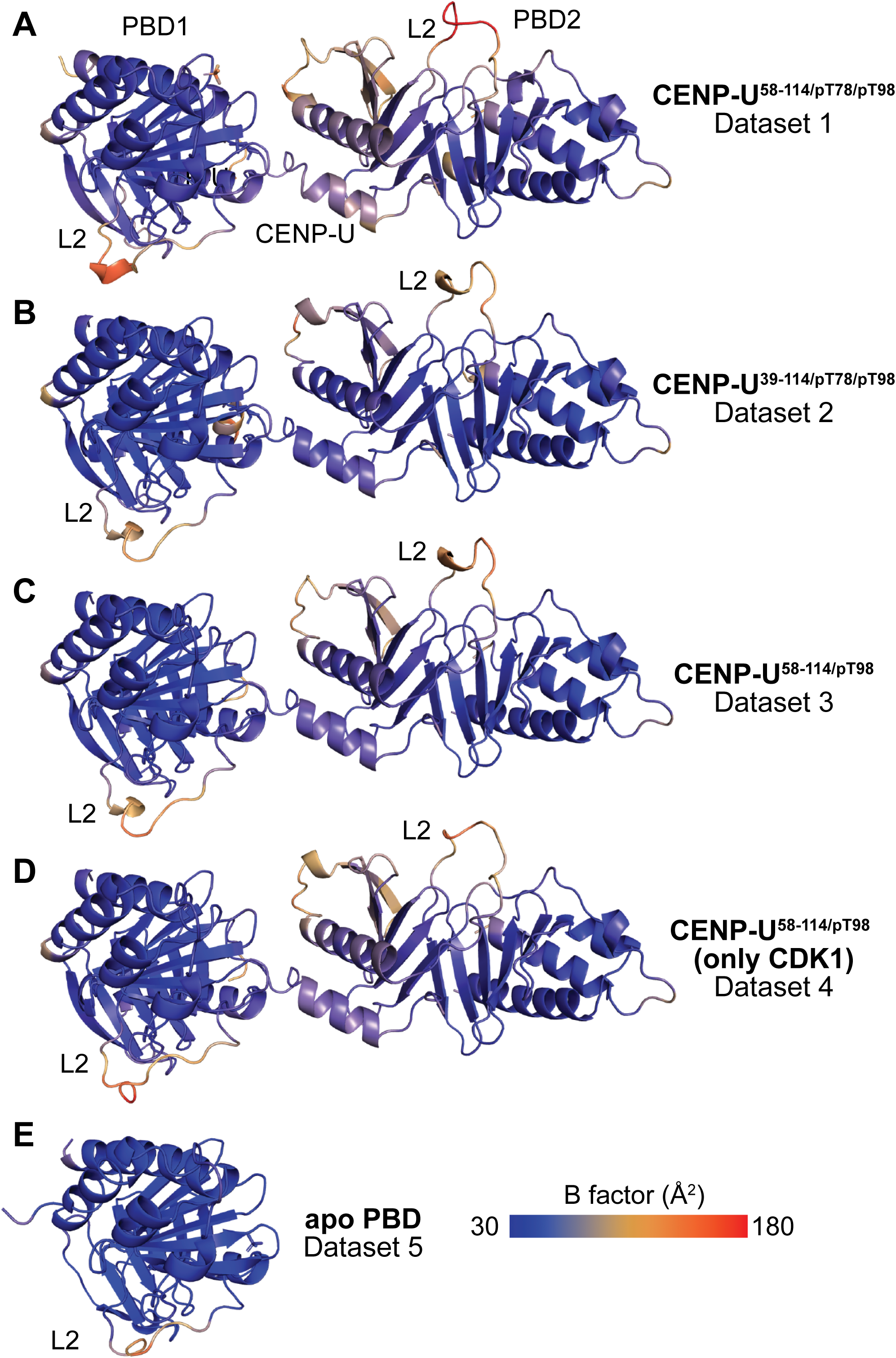
**(Associated with Figures 5 and 6)** (**A**-**E**) Distribution of atomic B-factors on the indicated experimental models. L2 loops in PBD1 and PBD2 are among the most mobile regions of the structures.

**Figure S9.**
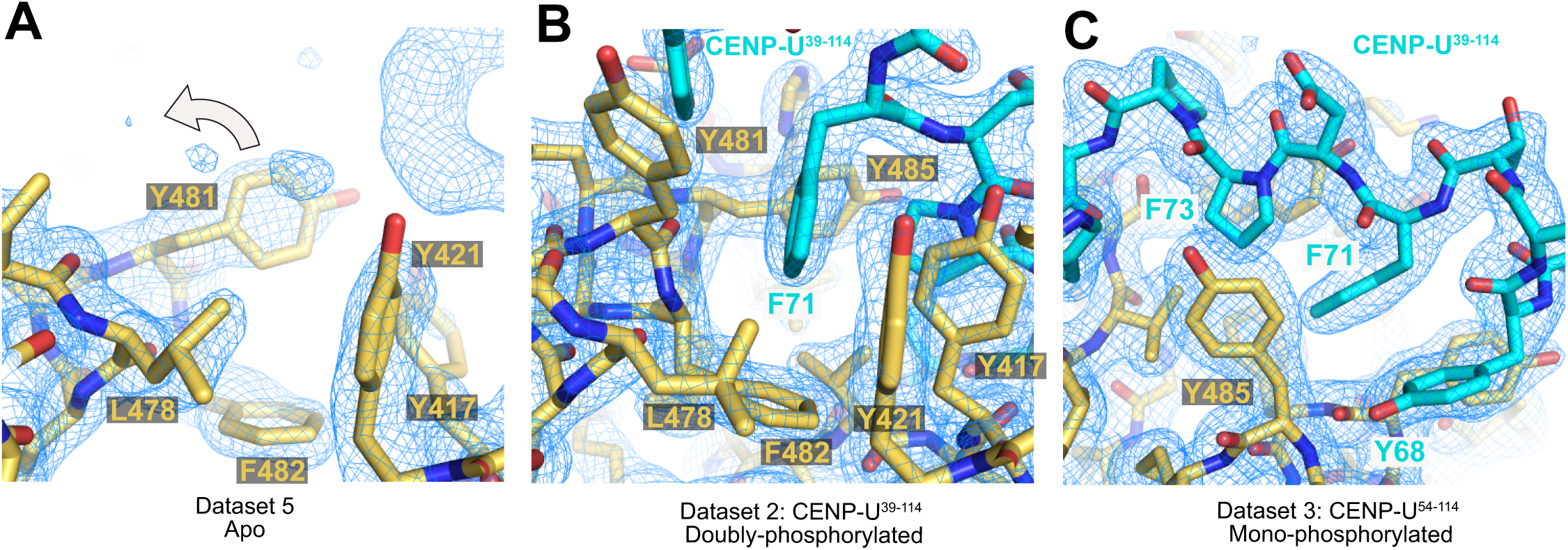
**(Associated with Figure 6)** (**A**) Position of the side chain of Tyr481 at the mouth of the cryptic pocket in the structure of the PBD calculated from Dataset 5 in absence of bound peptide. The arrow indicates the movement of the side chain of Tyr481 required to open the pocket and allow its occupation. (**B**) The side chain of Phe71 in Motif 1 in the cryptic pocket. The new position of Tyr481 is indicated. There is excellent electron density documenting the structural change. (**C**) A view of the cryptic pocket after an approximately 180° rotation from the view in panel B.

**Figure S10.**
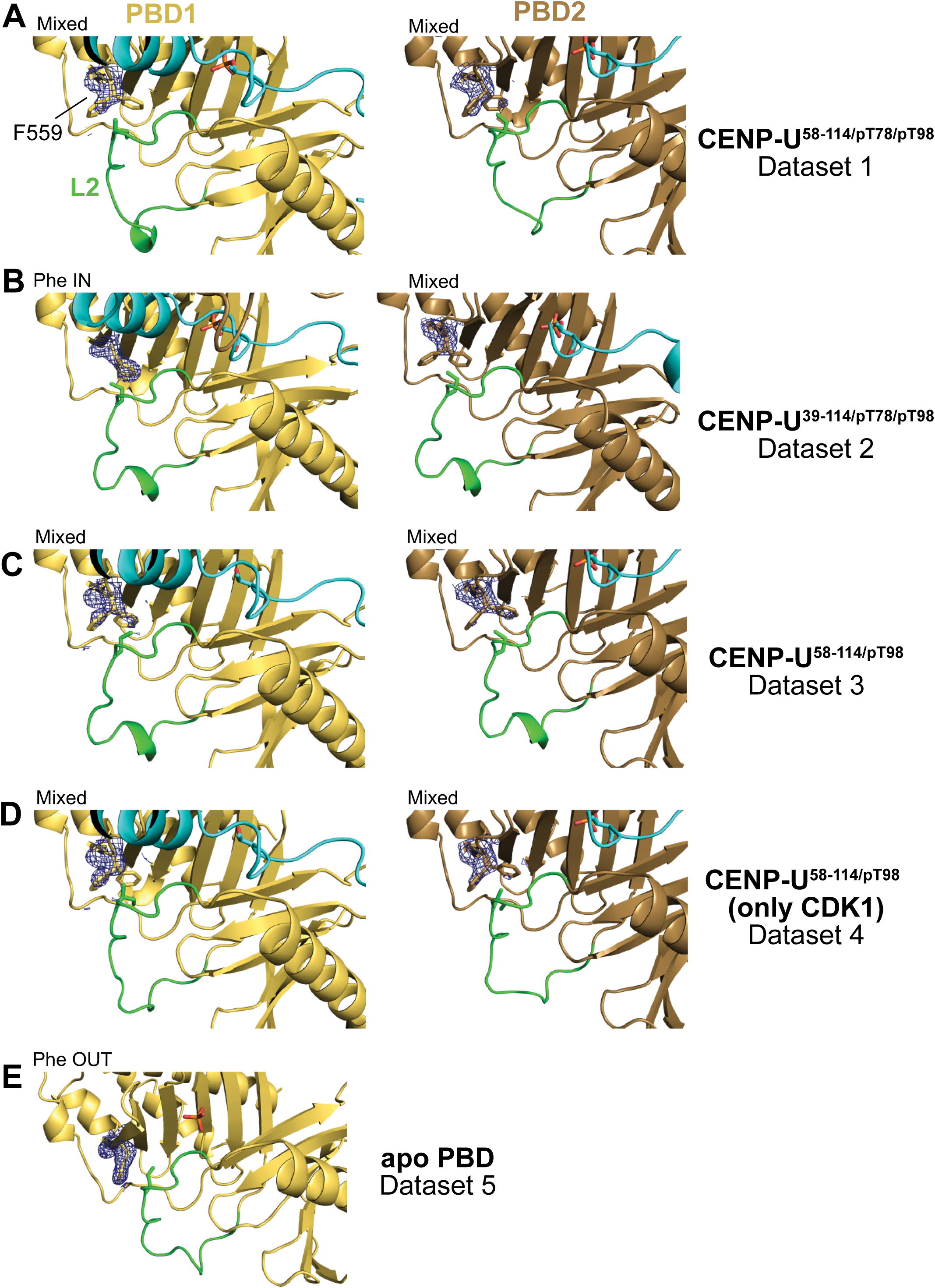
**(Associated with Figure 6)** (**A**-**E**) Electron density and modelling of the side chain of Phe599 (F599) in PBD1 and PBD2 of the indicated datasets. “Mixed” means that each of the two conformations, Phe IN and Phe OUT, is represented, to various levels, in the electron density, even if one of the forms predominates. “Pure” Phe IN or Phe OUT conformation are only observed in PBD1 in Dataset 2 and apo PBD in Dataset 5.

